# Astrocytic and endothelial GLUT1 restoration improves brain glucose homeostasis in GLUT1 deficiency syndrome

**DOI:** 10.64898/2026.03.04.709430

**Authors:** Shoko Tamura, Hiroko Shimbo, Nobuyuki Aruga, Mai Kawaguchi, Haruo Okado, Yuga Yasuda, Yuichiro Miyaoka, Ayano Sasakura, Kinya Matsuo, Hideaki Nishihara, Erika Seki, Kazunari Sekiyama, Kenichi Oshima, Shinobu Hirai

## Abstract

Glucose transporter 1 deficiency syndrome (GLUT1-DS) is a severe neurometabolic disorder caused by impaired brain glucose transport. Because GLUT1 has been classically viewed as predominantly localized to blood–brain barrier endothelial cells, gene therapy strategies have largely focused on endothelial GLUT1 restoration. However, whether this endothelial-centered model fully explains disease pathogenesis and therapeutic rescue has remained unclear. Using enhanced detection in mouse brain and human postmortem tissue, we show that GLUT1 is broadly expressed in astrocytes as well as endothelial cells, extending beyond perivascular endfeet into astrocytic somata and processes. Conditional Glut1 haploinsufficiency in either astrocytes or endothelial cells reproduced GLUT1-DS-like reductions in cerebrospinal fluid glucose availability and brain parenchymal glucose responses, together with cognitive and motor impairments. To determine the therapeutic consequence of this revised cellular model, we used complementary endothelial- and astrocyte-directed AAV delivery to restore GLUT1 in systemic Glut1 haploinsufficient mice. Single-compartment restoration produced partial benefit, whereas dual-compartment restoration more robustly improved cerebrospinal fluid glucose availability and cognitive function. These findings indicate that endothelial GLUT1 restoration alone is insufficient for robust rescue and redefine GLUT1-DS as a multicellular disorder of the neurovascular unit.

## Introduction

The human brain, though comprising only ∼2% of body mass, accounts for nearly 20% of systemic glucose utilization^1,2^. Glucose uptake in the brain is primarily mediated by facilitative glucose transporters (GLUTs), of which 14 isoforms exist in humans^3^. Among these, glucose transporter 1 (GLUT1) is the dominant isoform at the blood-brain barrier (BBB), expressed abundantly in vascular endothelial cells^4,5^.

Reduced expression or function of GLUT1 causes GLUT1 deficiency syndrome (GLUT1-DS), a rare metabolic encephalopathy first described in 1991^6^. Heterozygous *SLC2A1* mutations underlie most cases^7^, with an incidence estimated at 1 in 24,300-60,600 births^8,9^. Patients present with a broad spectrum of symptoms, including seizures, abnormal eye-head movements, cognitive and motor delay, and impaired head growth^10^. This clinical heterogeneity complicates the diagnosis of GLUT1-DS, and the true prevalence is likely underestimated. A reduced cerebrospinal fluid (CSF)-to-blood glucose ratio is the key diagnostic hallmark, seen in nearly all patients^11,12^.

Current therapy relies on ketogenic diets, but their efficacy for seizures and cognitive and motor dysfunction is only partial^10,13,14^. Safety concerns remain, particularly during pregnancy^15^. These limitations underscore the urgent need for curative approaches. Gene therapy using adeno-associated virus (AAV) vectors has emerged as a promising strategy, with proof-of-concept studies demonstrating benefit in animal models^16–19^. Endothelial cells (ECs) have been considered the primary therapeutic target. This is based on the fact that GLUT1 expression has been classically regarded as selective to the capillary endothelium^5,20^, and that Glut1 deletion restricted to endothelial cells phenocopies systemic haploinsufficiency^17,21^.

Nevertheless, evidence suggests that GLUT1 expression is not confined to ECs. Spatial transcriptomic datasets report Glut1 mRNA expression in microglia, oligodendrocytes, astrocytes, and neurons, although the corresponding protein distribution remains unclear (https://www.proteinatlas.org/). Several studies described astrocytic GLUT1 protein^22–24^, but these findings were inconsistent and difficult to interpret given the limited resolution of early staining approaches.

Here, we tested the hypothesis that GLUT1-DS is not solely an endothelial transport disorder but a multicellular disorder of the neurovascular unit involving both endothelial and astrocytic GLUT1. Using enhanced GLUT1 detection in mouse and human brains, together with cell-type-specific Glut1 haploinsufficient mouse models, we first determined whether astrocytic GLUT1 is broadly expressed and whether astrocytic, in addition to endothelial, GLUT1 is functionally required for maintaining brain glucose homeostasis. We then used systemic AAV-mediated compartment-selective GLUT1 supplementation to ask whether endothelial-only restoration is sufficient for therapeutic rescue, or whether coordinated restoration across endothelial and astrocytic compartments provides superior recovery of brain glucose availability and neurological function. Finally, as supportive translational data, we explored candidate SLC2A1 regulatory elements that may help guide future strategies for endogenous-like GLUT1 expression control.

## Results

### GLUT1 is expressed in both endothelial cells and astrocytes in mouse and human brains

We first assessed the cellular distribution of Glut1 (GLUT1) in the mouse and human brain. In mice, Glut1 was abundantly expressed across the brain parenchyma (Fig. 1A). While conventional immunohistochemistry primarily localized Glut1 to vascular ECs, enhanced detection with the avidin-biotin complex (ABC) method revealed extensive expression in astrocytes as well. Notably, the labeling extended continuously from perivascular endfeet along the soma to distal astrocytic processes (Fig. 1B). Similarly, in the human postmortem brains, GLUT1 was strongly expressed in ECs and weakly but broadly in astrocytes (Fig. 1C). These findings indicate that astrocytic GLUT1 expression is not confined to perivascular regions but is distributed throughout the entire astrocytic soma and processes.

**Figure 1.**
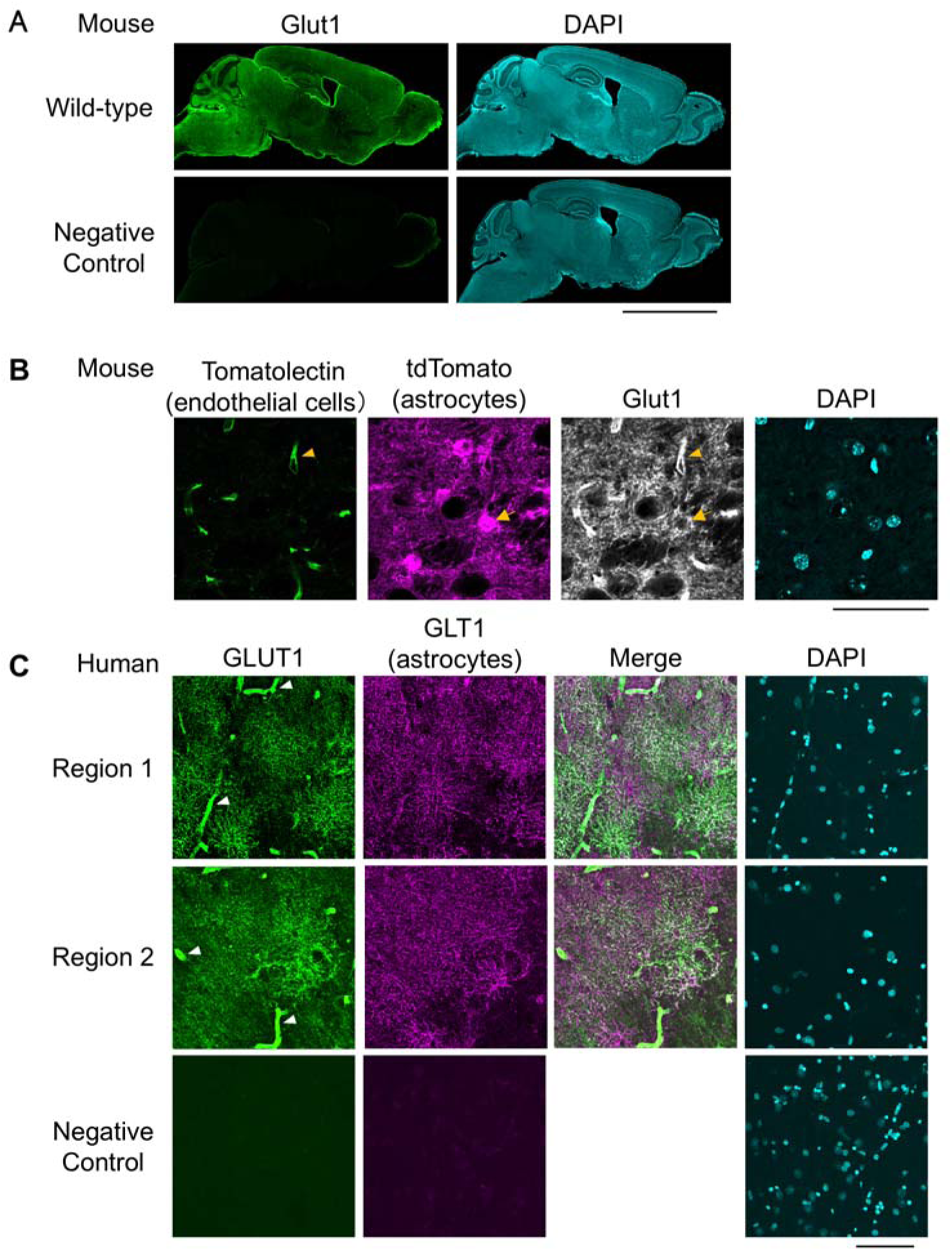
Glut1/GLUT1 expression in mouse and human brain. (A) Representative sagittal brain sections from wild-type mice immunostained for Glut1 (green) and counterstained with DAPI (cyan). Negative control sections were processed in parallel without the primary anti-Glut1 antibody. Scale bar, 5 mm. (B) High-magnification images of the thalamic ventral posteromedial nucleus (VPM) from 2-month-old Aldh1l1-CreERT2; Rosa26-LSL-tdTomato (flex tdTomato) mice. Endothelial cells were labeled with tomato lectin (green), astrocytes with tdTomato (magenta), Glut1 immunoreactivity is shown in grayscale, and nuclei are counterstained with DAPI (cyan). Yellow arrowheads indicate Glut1 signal associated with lectin-positive vessels, and yellow arrows indicate Glut1 signal in tdTomato-positive astrocyte somata. Scale bar, 50 µm. (C) Representative images from human hippocampus without neurological disease stained for GLUT1 (green) and the astrocytic marker GLT1 (magenta), with merged images and DAPI (cyan). White arrowheads indicate vascular GLUT1 staining. Region 1 and Region 2 correspond to the stratum oriens (or alveus) and the pyramidal cell layer, respectively. Negative control sections were processed without primary antibodies. Scale bar, 50 µm. Glut1/GLUT1 immunostaining was visualized using tyramide signal amplification (TSA).

### Both astrocytic and endothelial GLUT1 deficiency contribute to cognitive and motor impairments

To determine the functional relevance of astrocytic GLUT1, we compared systemic Glut1 heterozygous mice (Glut1 del/+) with conditional heterozygotes lacking one allele in astrocytes (Aldh1l1-Cre; Glut1 flox/+) or ECs (Tie2-Cre; Glut1 flox/+) (Fig. 2). Systemic Glut1 del/+ mice have been reported to show cognitive impairments resembling those in GLUT1 deficiency syndrome (GLUT1-DS) patients^21^. In both the object location test (OLT) and novel object recognition test (NORT), astrocyte-specific and EC-specific heterozygotes displayed significantly lower discrimination index scores than wild-type (WT) controls (OLT: p < 0.0001 and p = 0.0002; NORT: p = 0.0012 for both), closely paralleling deficits in Glut1 del/+ mice (Fig. 2A, B).

**Figure 2.**
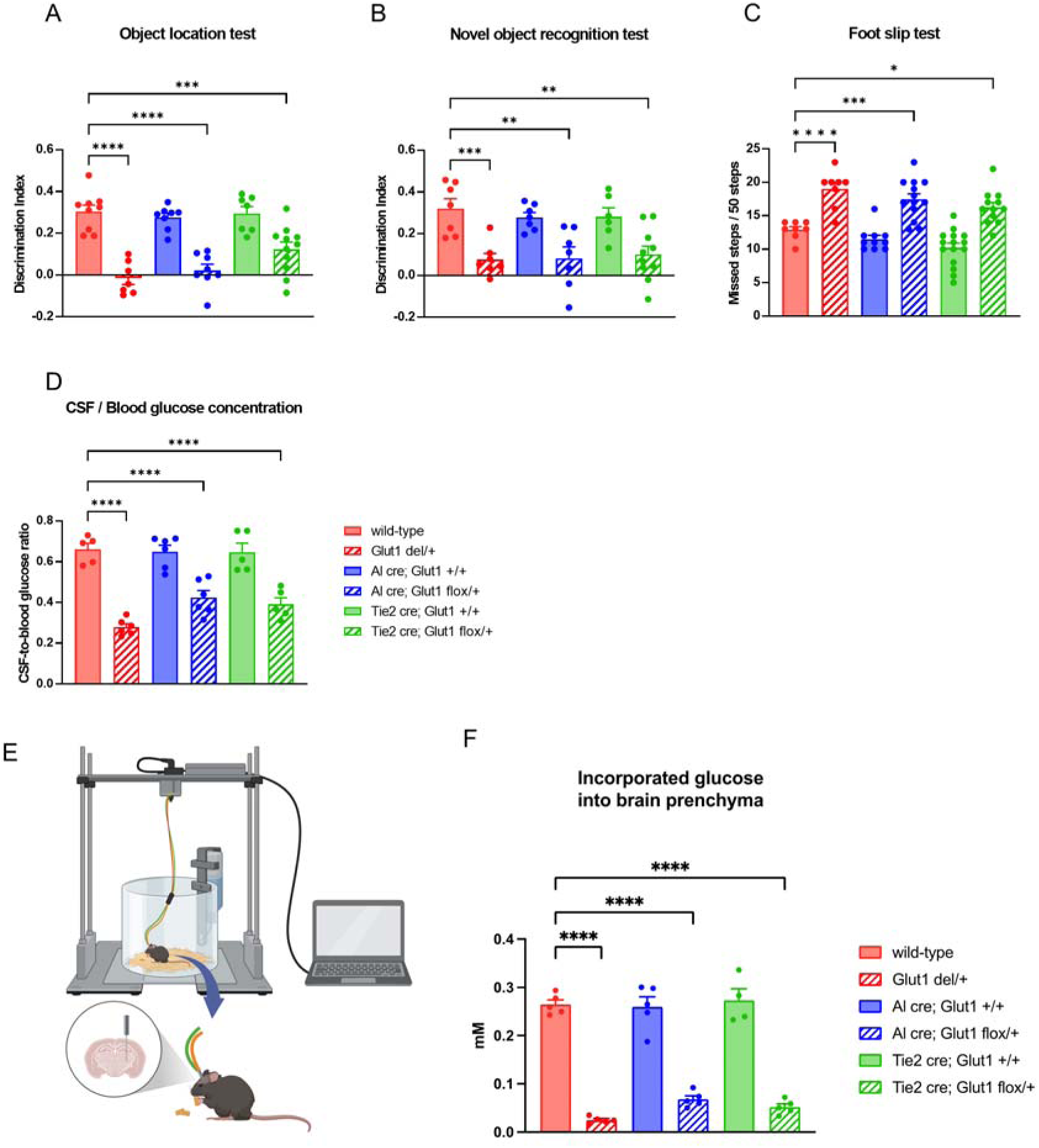
Astrocyte- and endothelial-specific Glut1 haploinsufficiency recapitulates key features of GLUT1 deficiency syndrome. (A-C) behavioral analyses in six groups: wild-type, systemic Glut1 del/+ mice, Aldh1l1-Cre (Al cre); Glut1 +/+ mice, Al cre; Glut1 flox/+ mice, Tie2-Cre; Glut1 +/+ mice, and Tie2-Cre; Glut1 flox/+ mice (n = 6-15 mice per group). (A) Object location test (OLT) and (B) novel object recognition test (NORT) were used to assess spatial and recognition memory (discrimination index). (C) The foot-slip test evaluated motor coordination (missed steps per 50 steps). (D) CSF-to-blood glucose ratio measured in the same cohorts (n = 5-6 mice per group). Blood glucose levels were comparable across groups (see Figure S1). (E) Schematic of the *in vivo* recording setup for thalamic interstitial fluid (ISF) glucose measurement. Illustration created with BioRender. (F) ISF glucose measured following oral glucose administration (0.03 g per mouse, gavage). The increase in ISF glucose was markedly blunted in systemic, astrocyte-specific, and endothelial-specific Glut1 haploinsufficient mice compared with wild-type controls (n = 4-5 mice per group). Data are presented as mean ± SEM; dots represent individual mice. Statistical significance was determined by one-way ANOVA followed by Dunnett’s multiple-comparison test (*p < 0.05, **p < 0.01, ***p < 0.001, ****p < 0.0001).

Motor coordination, one of the hallmark deficits in GLUT1-DS, was then evaluated using the slip test. As previously reported, EC-specific Glut1 deficiency impairs rotarod performance^17^. Consistent with this, EC-specific heterozygotes showed significant deficits compared with WT controls, and notably, astrocyte-specific heterozygotes exhibited comparable impairments (p < 0.0001, p = 0.0006, and p = 0.0180, respectively) (Fig. 2C). Together, these results indicate that both astrocytic and endothelial GLUT1 are indispensable for maintaining normal cognitive and motor functions.

### Glut1 deficiency in astrocytes and ECs lowers CSF-to-blood glucose ratio without altering blood glucose

Because a reduced CSF-to-blood glucose ratio is a diagnostic hallmark of GLUT1-DS, we next assessed glucose concentrations (Fig. 2D). While blood glucose levels were unchanged across groups (Figure S1), CSF-to-blood glucose ratio was significantly reduced in Glut1 del/+, Aldh1l1-Cre; Glut1 flox/+, and Tie2-Cre; Glut1 flox/+ mice compared to WT controls (p < 0.0001 for each). These results confirm that GLUT1 in both astrocytes and ECs is necessary for sustaining CSF glucose homeostasis.

### Both astrocytic and endothelial GLUT1 are required for brain glucose uptake

To directly evaluate brain glucose uptake, we monitored thalamic interstitial fluid (ISF) glucose concentrations after oral glucose administration (0.03 g per mouse) (Fig. 2E, F). Thalamic glucose uptake, as identified by [18F]-FDG positron emission tomography, is known to be reduced in GLUT1-DS patients^25^. ISF glucose elevation was markedly attenuated in systemic, astrocyte-specific, and EC-specific Glut1 heterozygotes compared with WT controls (p < 0.0001 for each). These findings establish that astrocytic and endothelial GLUT1 jointly enable glucose entry into the brain parenchyma in response to systemic glucose fluctuations.

### Astrocyte-directed AAV delivery enables testing of dual-compartment GLUT1 restoration

To determine whether astrocytic GLUT1 supplementation contributes to therapeutic rescue in vivo, we required an AAV vector capable of preferential gene delivery to astrocytes after systemic administration. Because conventional systemic AAV vectors show limited or nonselective astrocyte transduction in the CNS, we selected candidate capsids by reanalyzing previously published AAV insertion library datasets. This analysis identified heptapeptide motifs associated with astrocyte/CNS tropism. Based on motif enrichment and branch-aware prioritization, three candidate variants were selected for experimental validation and designated AAV-AST0, AAV-AST1, and AAV-AST3 (Figure S2A, B). These variants shared a conserved T(L/A/T)xxPF[K/L] core architecture while retaining branch-specific sequence differences.

To functionally validate these candidates, we performed intravenous administration in mice. AAV-AST0 failed to traverse the murine BBB and showed only limited CNS transduction (Figure S2). In contrast, AAV-AST3 crossed the BBB and robustly transduced astrocytes, but it also transduced neurons at a similar magnitude, prompting us to exclude it from further analyses (Figure S2). Among the variants tested, AAV-AST1, carrying the TALKPFL insertion, displayed the most favorable profile, combining efficient BBB penetration with pronounced astrocyte selectivity. We therefore selected AAV-AST1 as the lead candidate and hereafter refer to it as AAV-AST.

Following intravenous delivery, AAV-AST mediated widespread and efficient gene transfer throughout the brain, with a strong preference for astrocytes and minimal transduction of neurons and other CNS cell types (Fig. 3A). Astrocyte tropism was particularly pronounced in juvenile mice (Fig. 3B, C). We next evaluated transduction in primary human astrocytes. Compared with the control capsids AAV9 and PHP.eB, AAV-AST achieved significantly higher transduction efficiency (Fig. 3D-F). In mice, PHP.eB has been shown to cross the BBB efficiently and mediate broad CNS transduction, including astrocytes^26^. AAV9 is an AAV serotype currently used for intravenous gene therapy in human neurological diseases^27^. Together, these results indicate that AAV-AST provides sufficient astrocyte-directed delivery to test whether astrocytic GLUT1 supplementation contributes to therapeutic rescue *in vivo*.

**Figure 3.**
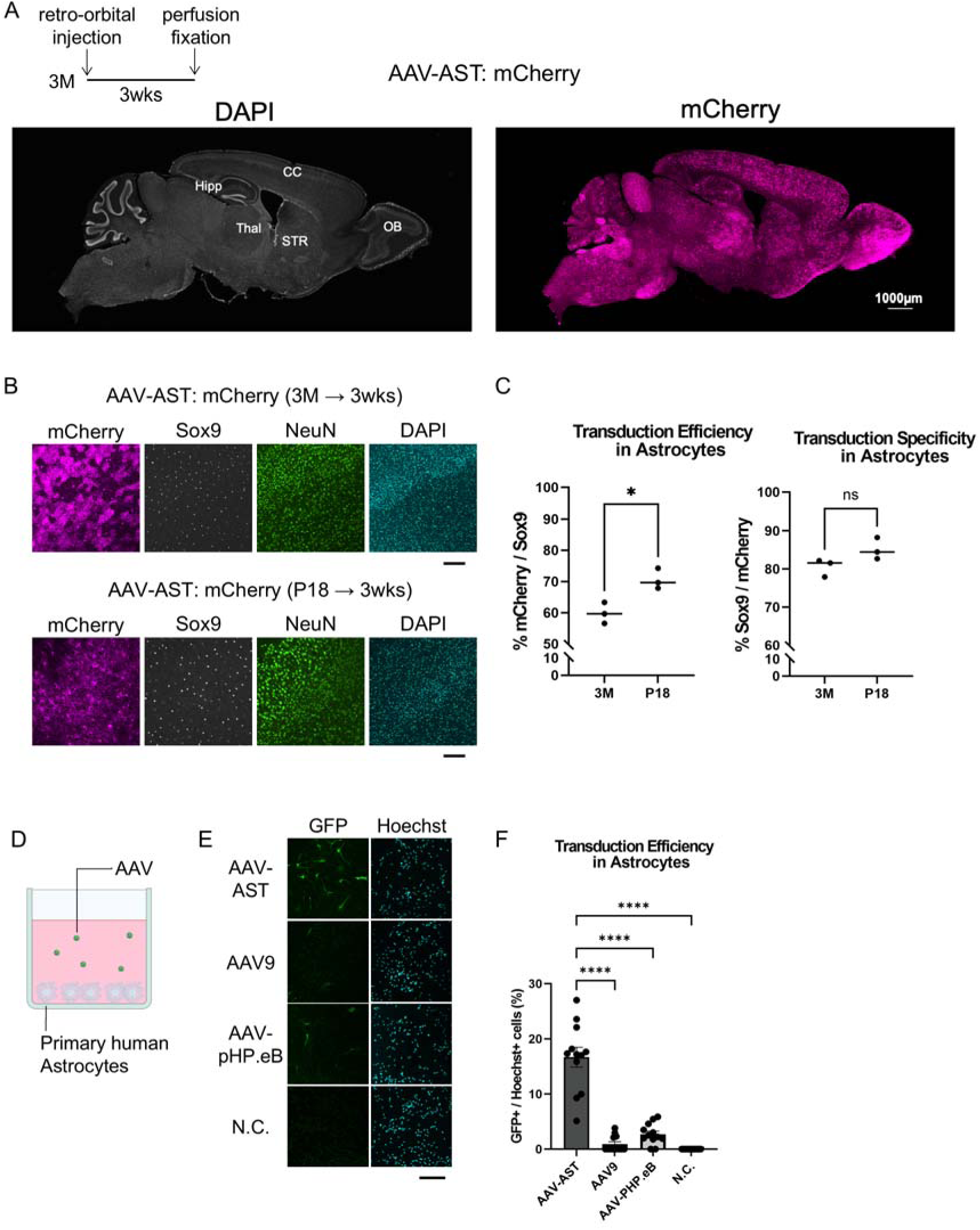
Astrocyte-selective transduction of AAV-AST in vivo and in vitro. (A) Representative sagittal brain sections showing mCherry expression (magenta) after retro-orbital administration of AAV-AST-CAG-mCherry into adult mice (3 months; 100 µL, 3.3 × 10^11^ vector genomes (vg) per mouse). DAPI is shown in gray. Labeled regions: OB, olfactory bulb; CC, cerebral cortex; Thal, thalamus; STR, striatum; Hipp, hippocampus. Scale bar, 1,000 µm. (B) Representative immunofluorescence images of the somatosensory cortex 3 weeks after AAV administration at postnatal day 18 (P18) or 3 months (3M). mCherry (magenta), SOX9 (astrocytes, gray), NeuN (neurons, green), and DAPI (nuclei, cyan). Scale bar, 200 µm. (C) Quantification of astrocyte transduction efficiency (left; % of SOX9^+^ cells expressing mCherry) and astrocyte specificity (right; % of mCherry^+^ cells that are SOX9^+^). Data are mean ± SEM (n = 3 mice per group). Two-sided Welch’s t-test; *p < 0.05; ns, not significant. (D) Schematic of the in vitro astrocyte transduction assay using AAVs packaging a CAG-driven GFP reporter (CAG-GFP). Primary human astrocytes were cultured in monoculture, transduced with AAVs, and fixed 3 days later for fluorescence imaging. Created with BioRender.com. (E) Representative fluorescence images of astrocytes transduced with the indicated AAVs (GFP, green; Hoechst, cyan). N.C., non-transduced control. Scale bar, 200 µm. (F) Quantification of astrocyte transduction efficiency corresponding to (E) (GFP^+^/Hoechst^+^ cells, %). Data are mean ± SEM. Statistical significance was determined by one-way ANOVA followed by Tukey’s multiple-comparison test (****p < 0.0001).

### Dual endothelial and astrocytic GLUT1 restoration provides superior rescue over single-compartment targeting

To determine the extent to which dual targeting is required for phenotypic rescue, we performed intravenous rescue experiments in Glut1 del/+ mice using AAV vectors that preferentially transduce vascular endothelial cells (AAV-BR1^28^ and AAV-X1.1^29^) and an astrocyte-tropic vector (AAV-AST). Vectors encoding GLUT1-myc-T2A-mCherry under the CAG promoter were administered by retro-orbital injection between postnatal days 15 and 19 (P15-19) (Figure S3), either to restore GLUT1 in a single cell type (endothelium or astrocytes) or simultaneously in both. In behavioral analyses, object location memory (OLT) was significantly improved by either endothelial- or astrocyte-directed supplementation, whereas the dual-targeting regimen produced the most pronounced rescue relative to vehicle-treated heterozygotes (Fig. 4A). Similarly, the CSF-to-blood glucose ratio was partially restored by single-cell-type supplementation and further enhanced by dual targeting, approaching wild-type levels (Fig. 4E). In contrast, in the novel object recognition test (NORT), single-cell-type supplementation did not produce a detectable rescue, whereas dual supplementation restored performance, indicating that concurrent restoration in both compartments is required for this cognitive domain (Fig. 4B). Motor coordination deficits assessed by the foot-slip test were not significantly improved in any rescue group compared with Glut1 del/+ controls under this treatment paradigm (Fig. 4C). Random-fed blood glucose remained unchanged across all viral and control groups (Fig. 4D). These results indicate that endothelial GLUT1 restoration provides partial therapeutic benefit but is insufficient to restore all glucose-related and cognitive endpoints. The additional rescue achieved by astrocytic supplementation supports the concept that GLUT1-DS therapy should target the multicellular glucose-transport architecture of the neurovascular unit rather than the endothelial compartment alone.

**Figure 4.**
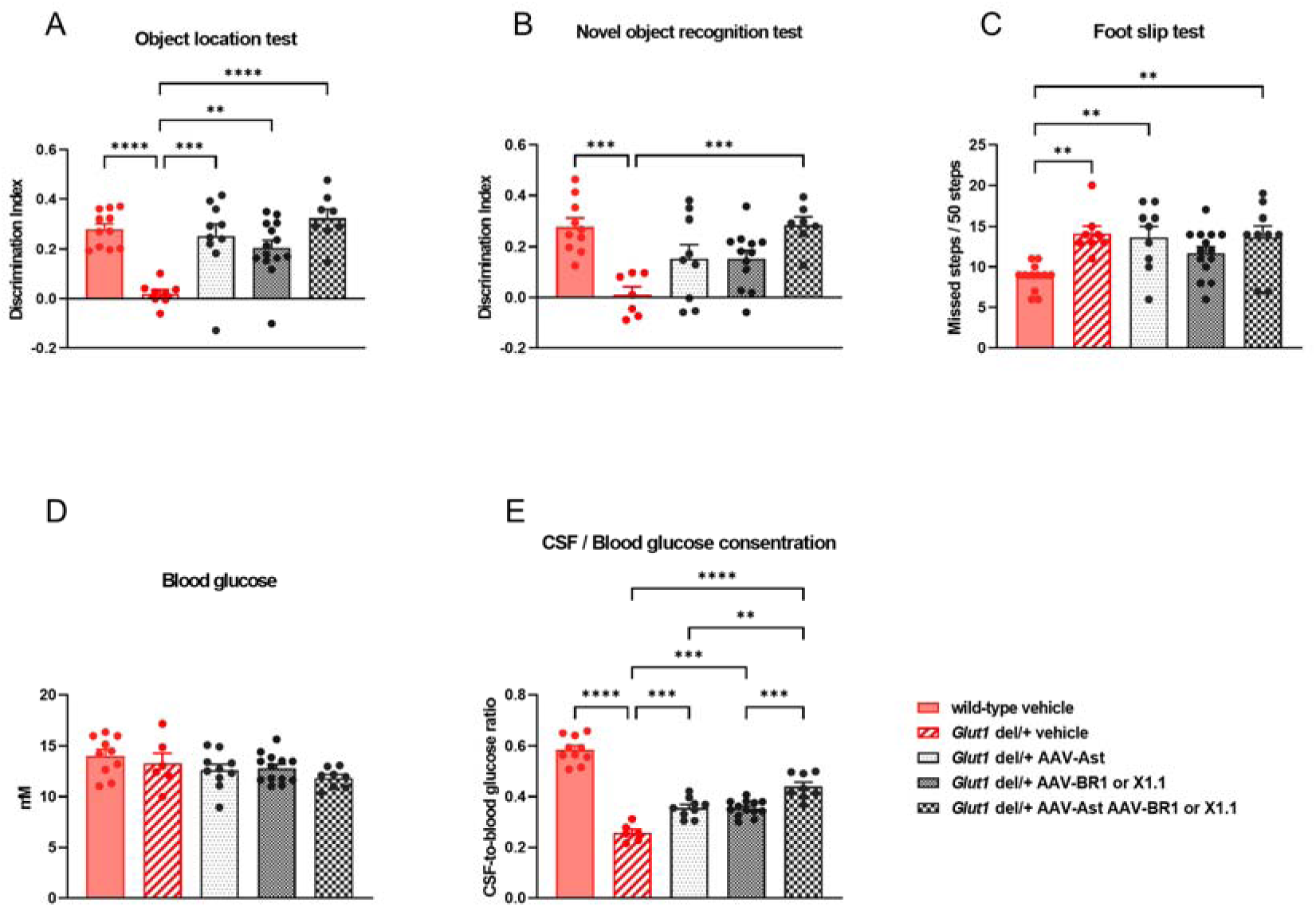
Cell-type–targeted AAV-Glut1 supplementation differentially rescues GLUT1-DS–like phenotypes. (A-C) Behavioral analyses in five groups: wild-type + vehicle, Glut1 del/+ + vehicle, Glut1 del/+ + AAV-AST-Glut1 (astrocyte-targeted), Glut1 del/+ + AAV-BR1 (or X1.1)-Glut1 (endothelial-targeted), and Glut1 del/+ + AAV-AST-Glut1 + AAV-BR1 (or X1.1)-Glut1. AAVs were administered between postnatal days 15 and 19 (P15-19) via retro-orbital injection, and behavioral testing was performed starting at P48 (n = 5-14 mice per group). (A) Object location test (OLT) and (B) novel object recognition test (NORT) were used to assess memory performance (discrimination index). (C) Foot-slip test evaluated motor coordination (missed steps per 50 steps). (D, E) Blood glucose concentration (D) and CSF-to-blood glucose ratio (E) were measured (n = 6-14 mice per group). Data are presented as mean ± SEM; dots represent individual mice. Statistical significance was determined by one-way ANOVA followed by Tukey’s multiple-comparisons test (****p < 0.0001, ***p < 0.001, **p < 0.01, *p < 0.05).

### Candidate SLC2A1 regulatory elements support endogenous-like GLUT1 expression control

Having established that optimal phenotypic rescue requires restoring GLUT1 in both endothelial cells and astrocytes, we next asked whether therapeutic expression could be placed under endogenous-like transcriptional control. We therefore prioritized candidate cis-regulatory elements at the SLC2A1 (GLUT1) locus by integrating H3K27ac ChIP-seq tracks in the UCSC Genome Browser (https://genome.ucsc.edu/index.html), ENCODE-annotated candidate regions, and a previously reported SLC2A1 regulatory sequence, which was used as a reference sequence in subsequent analyses^30^ (Fig. 5A). Nine candidate sequences (a-i) were cloned upstream of an mCherry reporter and co-transfected with a CAG-GFP plasmid as a transfection control (Fig. 5B, C and Figure S4). In both human brain microvascular endothelial cells (hBMECs) (Fig. 5B, Figure S4A) and primary cultured human astrocytes (Fig. 5C and Figure S4B), only Region d exhibited reporter activity significantly higher than Ref. when quantified after normalization to CAG-GFP. These results identify Region d as a candidate cis-regulatory element for driving SLC2A1-like expression and support its potential use in combination with endothelial- and astrocyte-tropic AAV vectors for dual-targeted GLUT1 supplementation.

**Figure 5.**
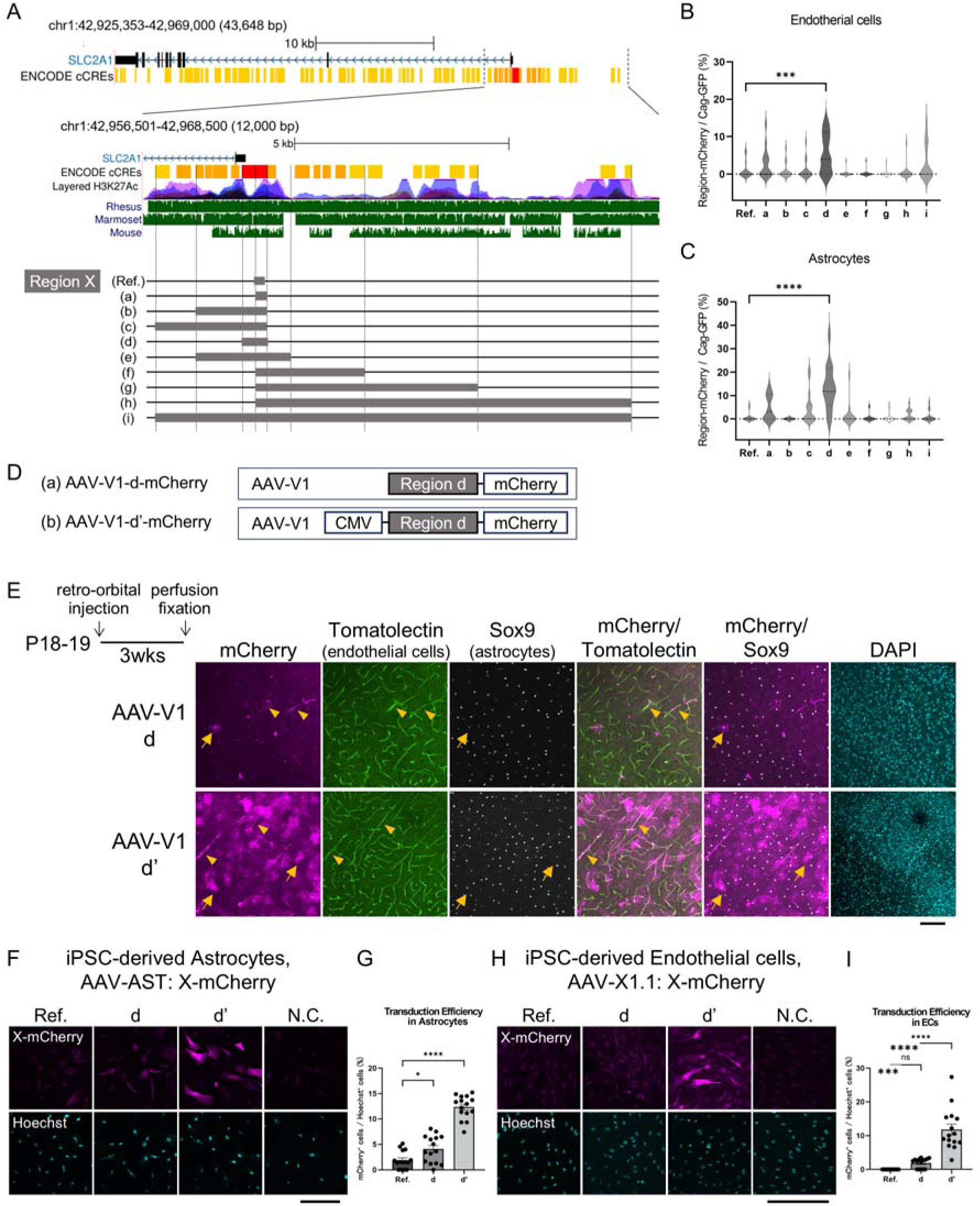
Candidate SLC2A1 regulatory elements support endogenous-like GLUT1 expression control. **(A)** UCSC Genome Browser view of the human *SLC2A1* locus (GRCh38/hg38), showing ENCODE candidate cis-regulatory elements (cCREs; orange), layered H3K27ac enrichment (purple), and sequence conservation across rhesus macaque, marmoset, and mouse (green). Gray boxes indicate the candidate regulatory fragments, including Ref. and Regions a–i, selected for functional testing and cloned upstream of an mCherry reporter. **(B, C)** Quantification of reporter activity in hBMECs **(B)** and primary cultured human astrocytes **(C)**. Reporter activity was quantified as the percentage of mCherry-positive cells among CAG-GFP-positive cells. Experiments were performed as shown in Supplementary Fig. 4A and 4B, respectively. n = 3 independent experiments; 15 fields per experiment. Violin plots show the median and interquartile range. Statistical significance versus Ref. was assessed by one-way ANOVA followed by Dunnett’s multiple-comparisons test; ***P < 0.001, ****P < 0.0001. **(D)** Schematic of AAV reporter constructs containing Region d, with or without an upstream CMV enhancer element, packaged into AAV-V1 capsids for in vivo analysis. **(E)** Representative confocal images of cortical sections from wild-type mice after retro-orbital administration of the indicated AAV-V1 vectors. mCherry is shown in magenta, tomato lectin-labeled vascular endothelial cells in green, SOX9-positive astrocytes in white, and DAPI in cyan. Region d drove reporter expression predominantly in vascular endothelial cells, with detectable expression in astrocytes. Addition of an upstream CMV enhancer to Region d, termed Region dlZ, increased reporter signal without an obvious change in cell-type distribution. Yellow arrowheads indicate representative vascular endothelial cells, and yellow arrows indicate astrocytes. Scale bar, 100 µm. **(F, H)** Representative fluorescence images of human iPSC-derived astrocytes **(F)** and endothelial cells **(H)** transduced with AAV vectors carrying the indicated regulatory regions, Ref., d, or dlZ. AAV-AST vectors were used for astrocytes, and AAV-X1.1 vectors were used for endothelial cells. mCherry is shown in magenta and Hoechst in cyan. N.C., non-transduced control. Scale bars, 300 µm. **(G, I)** Quantification of reporter expression in human iPSC-derived astrocytes **(G)** and endothelial cells **(I)**. Reporter-positive cells were quantified as the percentage of mCherry-positive cells among Hoechst-positive cells. Data are shown as mean ± SEM. Statistical significance in **(G)** was determined by one-way ANOVA followed by Dunnett’s multiple-comparisons test, whereas **(I)** was analyzed using Welch’s t-tests followed by Holm correction; *P < 0.05, ***P < 0.001, ****P < 0.0001.

To evaluate whether the regulatory activity of Region d identified in vitro is preserved in vivo, a reporter construct containing this regulatory element was packaged into an AAV-V1 capsid and systemically administered to mice by retro-orbital injection (Fig. 5D). To interrogate the function of the regulatory element using a single capsid, we chose AAV-V1, an AAV serotype capable of transducing both vascular endothelial cells and astrocytes in mice^31^. Region d drove reporter expression primarily in vascular endothelial cells, with detectable expression in astrocytes, thereby partially recapitulating the endogenous SLC2A1 (GLUT1) expression pattern.

To enhance expression strength without altering cell-type-specificity, a CMV enhancer was placed upstream of Region d^32^ (hereafter referred to as Region dlZ; Fig. 5D). This modification increased reporter signal in both vascular endothelial cells and astrocytes without an obvious shift in cell-type distribution, suggesting that enhancer-coupled Region d can increase expression strength while retaining the overall endothelial- and astrocytic expression pattern observed with Region d alone (Fig. 5E).

Because direct validation in human tissues is not feasible, we next assessed the regulatory activity of these elements in human iPSC-derived cells using capsids selected for their distinct tropisms. Specifically, AST-Region Ref., AST-Region d and AST-Region dlZ, as well as X1.1-Region Ref., X1.1-Region d and X1.1-Region dlZ, were tested in human iPSC-derived astrocytes and endothelial cells, respectively (Fig. 5F-I). In these human cellular models, Region dlZ consistently drove the most robust reporter expression, indicating that the enhancer-coupled configuration retains regulatory activity in a human context. These experiments further provided a human cellular context in which to evaluate the activity of Region d and Region dlZ when delivered by astrocyte- or endothelial-directed AAV capsids.

Together, these findings identify Region d as a candidate SLC2A1 cis-regulatory element with activity in endothelial and astrocytic contexts, and Region dlZ as an enhancer-coupled derivative with increased reporter expression. By providing a strategy for more endogenous-like expression control across endothelial and astrocytic compartments, these regulatory modules help address a key translational challenge for future GLUT1 gene therapy. Thus, Region d and Region dlZ offer candidate regulatory components for implementing dual-compartment GLUT1 restoration with more physiological expression control.

## Discussion

In this study, we show that GLUT1 is expressed not only in vascular endothelial cells (ECs) but also widely in astrocytes in human and mouse brains. This distribution extends well beyond perivascular endfeet, indicating that astrocytes, long considered metabolic supporters, are themselves central to brain glucose homeostasis. Functionally, we demonstrate that reduction of Glut1 expression in either ECs or astrocytes alone was sufficient to induce phenotypes closely resembling GLUT1 deficiency syndrome (GLUT1-DS), including cognitive and motor impairments as well as reductions in cerebrospinal fluid (CSF)-to-blood glucose ratios.

In the conventional view, GLUT1-dependent glucose entry in GLUT1-DS has been attributed primarily to endothelial transport at the BBB, whereas astrocytes have been regarded mainly as downstream metabolic supporters of neuronal function^2,33,34^. Our findings support a revised model in which astrocytic GLUT1 is broadly distributed beyond perivascular endfeet and contributes to the maintenance of brain glucose availability, together with endothelial GLUT1. This multicellular view provides a mechanistic explanation for why endothelial-only GLUT1 restoration produces incomplete rescue and establishes dual endothelial and astrocytic restoration as a more rational therapeutic design principle for GLUT1-DS.

Previous spatial transcriptomic datasets have shown that Glut1 mRNA is expressed in multiple non-endothelial cell types, in addition to endothelial cells (https://www.proteinatlas.org/), **whereas protein-level detection by immunoblotting and immunohistochemistry has primarily identified GLUT1 in ECs**^5,20^. Astrocytic GLUT1 has been thought to localize mainly at perivascular endfeet^5,22^. By integrating these fragmented observations, our study demonstrates widespread GLUT1 distribution throughout astrocytic membranes in both human and mouse brains (Fig. 1). Interestingly, in mice, all genetically labeled astrocytes expressed GLUT1, whereas in human brains, some astrocytes expressed GLUT1 while others did not, consistent with the evolutionary expansion of astrocytic diversity^35,36^. These results suggest that human astrocytes may exhibit more specialized metabolic subtypes than those in rodents.

Systemic Glut1 heterozygous mice recapitulate cardinal features of GLUT1-DS, including reduced CSF-to-blood glucose ratios and impaired neurodevelopment, motor coordination, and seizure susceptibility^37^. EC-specific Glut1 heterozygotes similarly display reduced CSF glucose and motor impairments^4,17^. Extending these findings, we show here that astrocyte-specific heterozygotes also develop cognitive and motor deficits with decreased CSF-to-blood glucose ratios, thereby highlighting that astrocytic Glut1 contributes as critically as endothelial Glut1 to GLUT1-DS pathogenesis (Fig. 2).

Mechanistically, astrocytes take up glucose **and metabolize it into lactate, thereby contributing to neuronal metabolism through the astrocyte–neuron lactate shuttle**^1,2,33,34^. On the other hand, glucose itself is present in the extracellular space and has been measured via microdialysis^38,39^. In the present study, systemic, EC-specific, and astrocyte-specific Glut1 heterozygous mice all showed reduced CSF-to-blood glucose ratios, and our biosensor-based analysis revealed significantly impaired glucose entry into the brain parenchyma after oral glucose administration (Fig. 2). Together with the broad membrane distribution of GLUT1 in astrocytes, these findings suggest that astrocytes are not limited to lactate-mediated metabolic support, but may also contribute directly to maintaining extracellular glucose availability. Because GLUT1 operates as a facilitated transporter, astrocytes may mediate glucose exchange according to local concentration gradients, coupling intracellular and extracellular glucose availability^40^. Furthermore, given the close communication and compositional similarity between CSF and brain interstitial fluid^41^, reduced CSF glucose may reflect not only impaired endothelial glucose entry into the brain, but also defective maintenance of parenchymal glucose availability resulting from reduced astrocytic GLUT1 (Figure S5).

From a therapeutic perspective, the ketogenic diet remains the standard of care for GLUT1-DS, but its efficacy for cognitive and motor impairments is limited. AAV-based gene therapy has therefore emerged as a promising strategy. Most existing therapeutic concepts have focused on restoring GLUT1 at the endothelial blood–brain barrier, consistent with the classical view that endothelial GLUT1 is the principal determinant of brain glucose entry. Our data support this concept in part, as endothelial-directed GLUT1 supplementation improved some endpoints. However, endothelial restoration alone did not fully normalize CSF glucose availability or cognitive function. In particular, novel object recognition was rescued only when astrocytic and endothelial compartments were targeted together. These findings indicate that endothelial replacement is beneficial but incomplete, and that effective GLUT1-DS gene therapy may require restoration of the multicellular glucose-transport system spanning both endothelial cells and astrocytes.

Notably, motor coordination was not rescued under our treatment paradigm, consistent with previous work showing that later interventions fail to restore motor function^19^. This aligns with evidence that motor phenotypes arise from early-life glucose insufficiency and that treatment efficacy is strongly dependent on the developmental stage at initiation^17,42,43^. Thus, earlier intervention may be necessary to fully rescue motor outcomes in GLUT1-DS, paralleling the high glucose demand of the developing human brain^44^. These findings further suggest that timing of intervention will be a critical variable in future therapeutic translation.

A key translational challenge for GLUT1-DS gene therapy is to reproduce the relevant cellular distribution of GLUT1 following systemic delivery. Our study identifies Region d as a candidate regulatory element that supports endogenous-like GLUT1 expression in both ECs and astrocytes (Fig. 5 and Figure S4). Enhancer-coupled Region dlZ increased reporter expression without an obvious shift in the overall endothelial and astrocytic expression pattern. Cross-context validation in human iPSC-derived astrocytes and endothelial cells further supports the relevance of this regulatory module for human translation. To implement dual-compartment restoration, complementary vector systems are required. X1.1 has been reported to efficiently transduce human vascular endothelial cells^29^, whereas AAV-AST provided astrocyte-directed delivery following systemic administration (Fig. 3 and Figure S2). In principle, combining endothelial- and astrocyte-directed AAV vectors with Region d or Region dlZ could provide a modular route toward endogenous-like GLUT1 restoration across the two cellular compartments disrupted in GLUT1-DS.

Together, our findings demonstrate that astrocytic and endothelial GLUT1 are both required to maintain brain glucose homeostasis and prevent key neurobehavioral phenotypes of GLUT1-DS. The central therapeutic implication is that GLUT1-DS gene therapy should not be designed solely as endothelial replacement at the BBB. Instead, effective restoration of brain glucose availability may require coordinated GLUT1 supplementation across endothelial and astrocytic compartments. This pathogenesis-guided principle provides a framework for next-generation therapeutic strategies for GLUT1-DS.

## Materials and methods

### Sex as a biological variable

Male and female mice were used in this study. Specifically, male mice were used for behavioral testing to maintain consistency across the multiple genetic models and therapeutic intervention experiments and to minimize potential variability associated with the estrous cycle. Mice of both sexes were used for non-behavioral assays. The study was not designed or powered to detect sex-dependent differences, and data from both sexes in the non-behavioral assays were analyzed together. Although the principal mechanisms identified in this study are expected to be relevant to both sexes, further studies are required to determine whether the magnitude of the behavioral phenotypes and therapeutic responses differs between male and female mice. Human postmortem tissues and human iPSC-derived cell lines were included as available. The study was not designed to evaluate sex-dependent differences in human samples or cell-based assays.

### Human postmortem brain tissue collection

Human postmortem brain tissues were obtained from the Tokyo Metropolitan Matsuzawa Hospital. The use of postmortem human brain tissues in the present study was approved by the Ethics Committee of Tokyo Metropolitan Institute of Medical Science (no. 23-26) and complied with the Declaration of Helsinki and its later amendments. All procedures were conducted with the informed written consent of the next of kin.

### Immunohistochemistry of Postmortem Brain Tissue

Tissue sections were permeabilized with PBS containing 0.3% Triton X-100 for 30 min at room temperature (R.T.), followed by quenching of endogenous peroxidase activity with 1% aqueous hydrogen peroxide for 20 min. Heat-induced antigen retrieval was performed in Tris-HCl buffer (pH 9.5) at 98°C for 1 hour. After blocking with 10% normal goat serum in PBS containing 0.3% Triton X-100 for 1 hour, sections were incubated overnight at 4°C with primary antibodies against GLUT1 (rabbit; 1:200, abcam, ab115730) and GLT-1 (guinea pig; 1:200, Millipore, AB1783).

Following three washes in PBS containing 0.3% Triton X-100, sections were incubated for 2 hours at RT with biotinylated goat anti-rabbit IgG (1:1000, Vector, BA-1000) to detect GLUT1. After another set of washes, signal amplification was performed using the VECTASTAIN Elite ABC Kit (PK-6100) for 30 min. Sections were then blocked with 5% normal donkey serum in PBS containing 0.3% Triton X-100 for 15 min, followed by TrueBlack® Lipofuscin Autofluorescence Quencher (5% in 70% ethanol, Cosmo Bio, Tokyo, Japan) treatment for 30 sec.

For fluorescence detection, GLUT1 signal was developed with TSA-Cy5 (1:500) for 10 min. Sections were then incubated with donkey anti–guinea pig IgG Cy3 (1:500, Merck Millipore, AP193C) to detect GLT-1, together with DAPI (1:100, Dojindo, D523) for 2 hours at R.T. After final washes, sections were mounted with fluorescence mounting medium.

### Animals

All experimental procedures were approved by the Animal Experimentation Ethics Committee of the Tokyo Metropolitan Institute of Medical Science (No. 24-034, approved on 1 April 2024). Animal care was provided in accordance with the institutional animal experimentation guidelines. All mice were maintained under controlled temperature (23lZ±lZ1lZ℃), humidity (50lZ±lZ10%), and 12:12 h light/dark cycle (lights on at 8:00 a.m.) with free access to a standard laboratory diet and water. All efforts were made to minimize the number of animals used and their suffering. Glut1 heterozygous mutant (Glut1 del/+) mice, used as a model for Glut1 deficiency syndrome (Glut1-DS)^37^, were generated by crossing CAG-Cre mice (a kind gift from Dr. Murakami^45^) with floxed Glut1 mice (The Jackson Laboratory, stock no. 031871). This cross results in Cre-mediated deletion of the floxed Glut1 gene in oocytes, leading to systemic Glut1 deficiency in the offspring. Mice harboring the Tie2-Cre (stock no. #004128), Aldh1l1-Cre (stock no. #023748), Aldh1l1-CreERT2 (stock no. #029655) drivers, tdTomato-floxed STOP reporter mice (tdTomato reporter mice) (stock no. 007914) and floxed Glut1 mice (stock no. #031871) were all obtained from the Jackson Laboratory, backcrossed over 10 generations to the C57BL/6J genetic strain background. Homozygous Glut1 flox/flox mice were mated with Tie2-Cre and Aldh1l1-Cre mice to generate Tie2-Cre; Glut1 flox/+ and Aldh1l1-Cre; Glut1 flox/+ conditional heterozygous mice, respectively.

### CSF and blood glucose measurements

CSF was extracted from the cisterna magna using a protocol described by Pegg et al.^46^. Mice were anesthetized by intraperitoneal injection of pentobarbital (34.5-51.8 mg/kg), which was chosen to minimize increases in blood glucose levels, and secured in the prone position. The head was tilted forward at an angle of approximately 25 degrees. Using a sterilized scalpel, the skin was incised, and the muscle layers were carefully separated to provide access to the cisterna magna, located between the occipital bone and the first cervical vertebra. CSF was collected from the cisterna magna using sterilized glass capillaries (1B150F-4, World Precision Instruments) connected to a winged needle and syringe assembly. Samples contaminated with blood were discarded. The glucose concentration in the collected CSF was measured using a blood glucose meter (Abbott Laboratories). Simultaneously with the CSF collection, blood was sampled from the tail tip. While the mouse was secured in a stereotaxic frame, a small incision was made at the tip of the tail using a razor blade, and 2-3 µL of blood was collected. The blood glucose level was immediately measured using a blood glucose meter (Abbott Laboratories).

### Measurement of glucose uptake into brain parenchyma

Extracellular glucose levels were measured using a commercially available in vivo biosensor recording system (Pinnacle Technology) according to the manufacturer’s instructions. A guide cannula was stereotaxically implanted to target the thalamus (AP −1.5 mm, ML −1.3 mm, DV −2.0 mm), and mice were allowed to recover for 1 week. Immediately before each recording, the glucose biosensor (standard mouse sensor) was calibrated using known glucose concentrations following the manufacturer’s protocol, and then inserted into the guide cannula. Recordings were initiated after the signal stabilized; mice were additionally allowed to equilibrate for 1 h after sensor insertion to ensure a stable baseline. Glucose (0.03 g per mouse) was administered by oral gavage, and extracellular glucose was continuously recorded at 1 Hz. Signals were transmitted from the potentiostat to a computer and acquired using Sirenia Acquisition Software (Pinnacle Technology). Glucose elevation was quantified as the difference between the baseline level immediately before gavage and the peak level observed after gavage. The glucose response was quantified as the peak ΔISF glucose (peak value minus the mean baseline value immediately before gavage).

### Behavioral tests

Prior to each behavioral test, mice were acclimated to the behavioral testing room for at least 30 minutes. The object location test (OLT) and novel object recognition test (NORT) were performed to assess memory function^47^, while motor function was evaluated using the slip test^48^.

Both the OLT and NORT consisted of three sessions: habituation, learning, and test. Mice were first placed in the center of the test apparatus (33 × 50 × 40 cm) and allowed to explore freely for 10 minutes, followed by a 1-2 hour return to their home cage. During the familiarization session, two identical objects (A and B) were placed in adjacent corners, 5 cm from the walls of the arena. Mice were reintroduced to the arena for 5 minutes, then returned to their home cage for 5 minutes. For the OLT test session, object B was moved to a diagonally opposite corner (designated B’), and mice were allowed to explore the arena for an additional 5 minutes. NORT was conducted immediately afterward. Mice were returned to the home cage for 5 minutes, and object A was replaced with a novel object (object C) in the same location. Mice were then placed back in the arena for another 5-minute test session.

Exploration times for objects A, B’, and C were recorded to calculate the discrimination indices as follows:

- OLT = (B’ − A) / (B’ + A)
- NORT = (C − B’) / (C + B’)
- where A, B’, and C represent the time spent exploring each object.

All behavioral tests were conducted under ambient lighting conditions of 10-15 lux. Mouse behavior was recorded using a ceiling-mounted charge-coupled device (CCD) camera positioned above the apparatus.

For the slip test, mice traversed a horizontal grid while being monitored with a side-mounted CCD camera. The number of times the hindlimb and forelimb falling through the grid was counted over 50 steps.

### Immunofluorescence staining of mouse brain tissue

Following transcardial perfusion with phosphate-buffered saline (PBS) and 4% paraformaldehyde (PFA), whole brains were extracted, post-fixed overnight at 4lZ°C, and cryoprotected in 20% sucrose at 4lZ°C for one week. Serial coronal sections (50lZμm thick) were prepared using a cryostat (CM3050 S; Leica Microsystems). Antigen retrieval was performed by incubating the sections in HistoVT One solution (Nacalai Tesque) at 70lZ°C for 30 minutes in a water bath. Sections were then permeabilized with a solution containing 0.2% Triton X-100, 1% Block Ace (DS Pharma Biomedical) in PBS for 30 minutes at room temperature. This was followed by overnight incubation with primary antibodies in PBS containing 0.4% Block Ace at room temperature. The immunostaining protocol continued with three washes in PBST (PBS with 0.05% Tween 20), followed by a 3-hour incubation with fluorophore-conjugated secondary antibodies diluted in PBS containing 0.4% Block Ace and 0.05% sodium azide. Nuclei were counterstained with 4lZ,6-diamidino-2-phenylindole (DAPI; Nacalai Tesque). Sections were subsequently washed three times in PBST, mounted in Fluoromount (Diagnostic BioSystems), and imaged using a FluoView FV3000 confocal laser scanning microscope (Olympus/Evident).

The following primary antibodies were used: goat anti-RFP (1:2000; Rockland, 200-101-379), guinea pig anti-Glut1 (1:1000; Frontier Institute, Af610), and/or rabbit anti-Sox9 (1:1000; Abcam, ab185966). Corresponding secondary antibodies included donkey anti-goat IgG conjugated to Cy™3 (1:1000; Jackson ImmunoResearch, 705-165-147), donkey anti-guinea pig IgG conjugated to biotin (1:1000; Jackson ImmunoResearch, 706-066-148), and/or donkey anti-rabbit IgG conjugated to Alexa™647 (1:1000; Jackson ImmunoResearch, 711-606-152). Tomato lectin (1:1000; Vector Laboratories, FL-1171) was used for vascular endothelial staining.

For enhanced immunostaining using horseradish peroxidase (HRP), endogenous peroxidase activity was quenched by incubating sections in 1% H₂O₂ in PBS for 20 minutes after antigen retrieval, followed by PBS washes. The standard immunostaining protocol was then followed. After the secondary antibody incubation and subsequent washes, sections were incubated with the avidin–biotin–peroxidase complex using the VECTASTAIN Elite ABC Kit (Vector Laboratories, PK-6100) for 30 minutes and washed three times with PBST. HRP activity was visualized using Tyramide Signal Amplification (TSA) with Cyanine5 (1:500; PerkinElmer, FP1171).

In Fig. 1, Aldh1l1-CreERT2; tdTomato reporter mice were administered tamoxifen intraperitoneally at 100lZmg/kg once daily for three consecutive days starting at P45. Brains were fixed and processed for immunostaining analysis three weeks after the final tamoxifen injection.

### Plasmid construction for AAV production

The AAV expression vector, pAAV-CAG-hGLUT1-MycTag-T2A-mCherry-WPRE, was constructed by custom DNA synthesis (VectorBuilder). The hGLUT1 coding sequence corresponds to NM_006516.4 (Table S1).

### AAV Production and Purification

pAAV-BR1 was generated by inserting the 7-amino acid peptide NRGTEWD between positions 588 and 589 of the AAV2 capsid^28^, along with an N587Q substitution (asparagine to glutamine). Similarly, pAAV-X1.1 was generated by inserting the 7-amino acid peptide DGAATKN between positions 588 and 589 of the AAV9 capsid^29,49^. The production of pAAV-AST is described in results.

HEK293T cells were cultured in standard medium (D-MEM supplemented with 10% heat-inactivated fetal bovine serum [hiFBS] and 1% penicillin–streptomycin [PS]) until reaching approximately 90% confluency. The cells were triple-transfected using polyethylenimine (PEI-Max) to produce AAV vectors, following the protocol described by Grieger et al^50^. (2006). Briefly, the transgene expression plasmid (pAAV-CAG-hGLUT1-MycTag-T2A-mCherry-WPRE), a packaging plasmid (pAAV-AST, pAAV-BR1, or pAAV-X1.1), and the pHelper plasmid were diluted and mixed in Opti-MEM (Life Technologies) to allow for the formation of DNA-PEI complexes.

Six to twelve hours after transfection, the culture medium was replaced with D-MEM containing 2% hiFBS and 1% PS. Viral particles were harvested from the medium 7-10 days post-transfection. For viral recovery, the collected supernatant was mixed with a 5× AAV precipitation buffer (200 g PEG 8000 [Sigma-Aldrich] and 73.05 g NaCl in 500 mL H_2_O) and incubated at 4 °C for 2-3 hours, as described by Guo et al^51^. The mixture was then centrifuged at 3,000 × g for 30 minutes at 4 °C. The resulting PEG precipitate was resuspended in Dulbecco’s PBS (D-PBS) and treated with SuperNuclease (Sino Biological) in the presence of 5 mM MgCl2 for 60 minutes to digest residual host cell DNA and RNA. The reaction was subsequently terminated by the addition of 6.5 mM EDTA.

Finally, the virus was purified using an iodixanol discontinuous gradient, as described by Zolotukhin et al^52^. Briefly, following centrifugation to remove cellular debris, the clarified supernatant was layered onto an iodixanol (OptiPrep) step gradient (15%, 25%, 40%, and 54%) in an Ultra-Clear tube (25 × 89 mm; Beckman). The gradient was ultracentrifuged at 28,000 rpm for 18 hours at 10 °C. Viral fractions were then collected from the bottom three-fourths of the 40% layer to the top one-fourth of the 54% layer (approximately 4-5 mL) using a 19G needle. The eluate was buffer-exchanged and concentrated from PBS to HBSS using a Vivaspin 100K centrifugal filter (Sartorius).

AAV purity was evaluated by SDS-PAGE, and viral genome titers were quantified by qPCR using the KAPA SYBR Fast qPCR Kit (Nippon Genetics). Primers targeting the WPRE sequence were used:

- Forward: 5lZ-CTGTTGGGCACTGACAATTC-3lZ
- Reverse: 5lZ-GAAGGGACGTAGCAGAAGGA-3lZ

The expression plasmid served as the standard for quantification.

### Peptide motif analysis

Selection of analyzed datasets: Four published AAV insertion library studies were analyzed^31,53–55^. From Kumar et al., motifs showing log₁₀ enrichment > 1 in GFAP-positive cells were extracted, clustered to remove near-identical variants, and representative sequences were selected. Together with five motifs from Nonnenmacher and one each from Kunze and Hanlon, these sequences comprised the reference motif set used to define the overall astrocyte/CNS-related motif landscape.

**Extraction and normalization:** Flanking residues derived from the VP3 587-590 loop (A-Q/A-Q) were trimmed, retaining only the 7-residue insertions between VP3 588-589 (pos1-pos7). The normalized 7-mer sequences were aligned to construct a position-frequency matrix (PFM) for visualization. Sequence logos were generated from the position-frequency matrix using a custom Python/Matplotlib script, with per-position letter heights scaled by information content.

**Branch-aware motif compatibility scoring:** Because several literature-derived astrocyte/CNS-related motifs converged on a limited set of closely related sequence branches, candidate prioritization was performed using a branch-aware motif compatibility score rather than a single pooled logo score. For candidate prioritization, four predefined PHP.B-like branches were used: TLAV-like, TALK-like, TTLK-like, and TLQx-like. The TLAV-like, TALK-like, TTLK-like, and TLQx-like branches were defined using representative motifs centered on TLAVPFK, TALKPFL, TTLKPFL/TTLKPFS, and TLQLPFK/TLQQPFK/TLQIPFK-related sequences, respectively. For each branch, a branch-specific position-frequency model was constructed from normalized 7-mer motifs using Dirichlet pseudocount smoothing (α = 0.1). Candidate sequences were scored against branch-specific position-frequency models using the mean log-likelihood across the seven positions, and the highest branch-specific value was retained as the branch-aware motif compatibility score. Scores were normalized to the top-ranked motif (= 1.0) for visualization and comparison.

**Selection for in vivo validation:** Candidate motifs ranked within the top-scoring group were prioritized for experimental testing. TLAVPFK was excluded because it is identical to the PHP.eB insertion sequence. TTLKPFL, TALKPFL, and TLQLPFK were therefore selected for in vivo validation.

### In vivo administration of AAVs

Wild-type C57BL/6J mice at postnatal day 18 (P18) or 3 months of age received retro-orbital injections of AAV-AST0, AAV-AST, or AAV-AST3 at 3 × 10^11^ viral genomes (vg) per mouse (Fig. 3A-C, Figure S2C, D). For in vivo evaluation of Region d regulatory activity (Fig. 5D, E), wild-type C57BL/6J mice at 3 months of age received retro-orbital injections of AAV-V1 reporter vectors carrying Region d or Region dlZ at a dose of 3 × 10^11^ vg per mouse. In both experiments, 3 weeks after administration, mice were transcardially perfused with PBS followed by 4% PFA, and brains were collected for immunofluorescence analysis as described above. Representative confocal images were acquired from the cerebral cortex under identical imaging settings across groups.

### Cell culture, transfection and AAV transduction

Human brain microvascular endothelial cells (hBMECs; ACBRI 376) and the specialized complete medium kit with serum and CultureBoost (4Z0-500-R) were purchased from Cell Systems. Human cortical astrocytes (HAs; Cat. #1800) and Astrocyte Medium (AM; Cat. #1801) were obtained from ScienCell Research Laboratories. All cells were maintained at 37lZ°C in a humidified incubator with 5% CO₂.

For hBMEC culture, coverslips were pretreated with the attachment solution supplied by Cell Systems (4Z0-201) according to the manufacturer’s instructions. Briefly, coverslips were incubated with the attachment solution, which was then recovered and reused. For human cortical astrocyte culture, plates or coverslips were coated overnight at 37lZ°C with poly-L-lysine (PLL; Sigma-Aldrich, P2636-100MG) at 1 μg/μL in Milli-Q water, washed three times with PBS and used for cell seeding.

For reporter assays in hBMECs (Fig. 5B, Figure S4A) and human cortical astrocytes (Fig. 5C and Figure S4B), cells were seeded in 24-well plates at densities of 20,000 cells per well and 22,800 cells per well, respectively, 24 h before transfection. Immediately before transfection, 250 μL of medium was removed from each hBMEC well. In astrocyte cultures, 250 μL of medium was temporarily collected and retained from each well before transfection. Cells were co-transfected with pAAV-hSLC2A1p-mCherry and pAAV-CAG-GFP at a 1:1 molar ratio, with a total of 500 ng DNA per well, using Lipofectamine LTX (Thermo Fisher Scientific, 15338100) according to the manufacturer’s instructions. Negative-control wells received no transfection treatment. At 6 h after transfection, 250 μL of medium was added to each hBMEC well, whereas the retained medium was returned to each astrocyte well. Cells were fixed the following day with 4% paraformaldehyde, and nuclei were counterstained with Hoechst. No immunostaining was performed; mCherry and GFP fluorescence were evaluated directly.

For AAV transduction assays in human cortical astrocytes (Fig. 3D-F), cells were seeded in 24-well plates and maintained in monoculture. The indicated AAV vectors carrying a CAG-driven GFP reporter were added at 1 × 10⁹ vg per well. Cells were fixed 3 d after transduction, counterstained with Hoechst and subjected to fluorescence imaging.

### Image acquisition and quantitative analysis

Fluorescence images were acquired using a BZ-X1000 microscope (Keyence) with a 10× objective lens and saved as 24-bit TIFF files. Quantitative image analysis was performed using custom Python scripts. For analysis, each fluorescence channel was processed separately using single-channel images converted to 8-bit grayscale prior to binarization, and all fixed thresholds were applied to 8-bit images (intensity range, 0-255). Within each experiment, identical analysis parameters were applied across all experimental groups. For each sample, five fields were randomly selected for analysis, excluding fields containing saturated signals or coverslip edges. Three independent experiments were performed on separate days for each condition. In assays using fixed thresholds, exposure time, gain, binning, and image size were kept constant within each assay. The analysis targets, binarization conditions, noise exclusion criteria, and quantitative metrics were defined for each experimental system as follows.

**Quantification of AAV transduction efficiency in human cortical astrocytes**: GFP and Hoechst 33342 (Nacalai Tesque, 04929-82) fluorescence images were analyzed. Hoechst images were binarized using the IsoData method, and objects smaller than 10 pixels were excluded as noise. GFP images were binarized using a fixed threshold of 48. For each nucleus, the overlap between the Hoechst-derived nuclear mask and the GFP signal was calculated, and nuclei with a GFP-overlapping area covering at least 30% of the nuclear area were classified as GFP-positive. Transduction efficiency was calculated for each field as the number of GFP-positive nuclei divided by the total number of Hoechst-positive nuclei.

**Quantification of hGLUT1 promoter activity in hBMECs**: GFP and mCherry fluorescence images were analyzed. Images were binarized using fixed thresholds (GFP, 28; mCherry, 49), and objects smaller than 30 pixels were excluded as noise. The number of valid objects detected after binarization was counted for each channel, and promoter activity was quantified for each field as the ratio of mCherry-positive objects to GFP-positive objects.

**Quantification of hGLUT1 assay (AAV transduction) in iPSC-derived astrocytes and hBMECs:** Quantification of hGLUT1 assay (AAV transduction) in iPSC-derived astrocytes and hBMECs: mCherry and Hoechst/DAPI fluorescence images were analyzed for both iPSC-derived astrocytes and hBMECs using the same image-processing workflow. Images were binarized using fixed thresholds for mCherry (18). For iPSC-derived astrocytes, Hoechst images were binarized using a fixed threshold (17), whereas for hBMECs, DAPI images were binarized using Otsu’s thresholding method. Nuclear objects smaller than 10 pixels were excluded as noise. For each nucleus, the overlap between the nuclear mask and the mCherry signal was calculated, and nuclei with an mCherry-overlapping area covering at least 30% of the nuclear area were classified as mCherry-positive. Expression efficiency was calculated for each field as the number of mCherry-positive nuclei divided by the total number of Hoechst/DAPI-positive nuclei.

**Quantification of hGLUT1 promoter activity in primary human astrocytes**: GFP and mCherry fluorescence images were analyzed. Images were binarized using fixed thresholds (GFP, 10; mCherry, 15), and objects smaller than 100 pixels were excluded as noise. The number of valid objects detected after binarization was counted for each channel, and promoter activity was quantified for each field as the ratio of mCherry-positive objects to GFP-positive objects.

### Construction of reporter plasmids

Reporter plasmids for subsequent recombinant adeno-associated virus (rAAV) production were constructed by fusing candidate promoter/enhancer elements to the mCherry reporter gene. Each expression cassette was flanked by inverted terminal repeat (ITR) sequences, and a woodchuck hepatitis virus posttranscriptional regulatory element (WPRE) was included to enhance transgene expression. Candidate cis-regulatory regions (regions a-i) were selected from the human SLC2A1 locus (chr1: 42,956,501-42,968,500; GRCh38/hg38) and amplified by PCR from human genomic DNA isolated from HEK293 cells. To generate reporter constructs, pAAV-CAG-mCherry-spacer-WPRE (Table S2) or pAAV-CAG-mCherry-WPRE (derived from pAAV-CAG-mCherry-spacer-WPRE by removal of the spacer sequence) was digested with XbaI and XhoI to produce a linearized backbone containing the mCherry reporter, WPRE, and ITR sequences. Promoter/enhancer fragments were then inserted upstream of mCherry within the ITR-flanked expression cassette using NEBuilder HiFi DNA Assembly Master Mix (New England Biolabs, E2621). Regulatory elements a, b, c, d, e and Ref. were cloned into pAAV-promoterless-mCherry-spacer-WPRE, whereas regulatory elements f, g, h, and i were cloned into pAAV-promoterless-mCherry-WPRE. For constructs containing regulatory elements a, b, c, e, f, and g, putative ATG sequences were removed by site-directed mutagenesis PCR after plasmid assembly. Region dlZ was generated by inserting a CMV enhancer sequence upstream of Region d into the XbaI-digested pAAV-promoterless-mCherry-WPRE backbone, yielding a construct in which the CMV enhancer-Region d tandem arrangement was positioned immediately upstream of mCherry (Table S3). Primer sequences and primer names are provided in Table S4.

The resulting plasmids were transformed into Escherichia coli DH5α competent cells (Takara Bio, 9057) and purified using a NucleoBond Xtra Midi kit (Takara Bio, 740410). A plasmid expressing GFP under the control of the CAG promoter was used as an internal control.

### Imaging and Statistical Analysis

Fluorescence observation and quantitative analysis were performed using an FV3000 confocal laser scanning microscope, with all images captured under identical laser power and detection settings across all samples. For quantification, images captured with a Keyence BZ-X1000 microscope were analyzed using ImageJ software (NIH). Reporter activity was defined as the percentage of mCherry-positive cells exceeding a fixed fluorescence intensity threshold among the total GFP-positive cell population. Quantification was performed using 15 fields per condition from 3 independent experiments; field-level measurements were averaged within each experiment, and n represents independent experiments.

### Administration of AAVs for Glut1 del/+ mice

AAVs were administered via the retro-orbital sinus^56^ to Glut1 del/+ mice between P15 and P19. Mice were anesthetized using a combination of three agents: medetomidine (0.375 mg/kg; Nippon Zenyaku Kogyo Co., Ltd., Fukushima, Japan), midazolam (2 mg/kg; Maruishi Pharmaceutical Co., Ltd., Osaka, Japan), and butorphanol (2.5 mg/kg; Meiji Seika Pharma Co., Ltd., Tokyo, Japan). Glut1 del/+ mice received AAV-BR1, AAV-X1.1 and/or AAV-AST at doses of 1.0-7.5lZ×lZ10¹¹ vg per mouse for each virus, diluted in 100lZμL of Hanks’ Balanced Salt Solution (HBSS) containing 0.5% Fast Green. Control littermates received 100lZμL of vehicle (HBSS with 0.5% Fast Green) via the same retro-orbital injection method.

### Culture of human iPSCs and generation of iPSC-derived astrocytes and EECM-derived brain microvascular endothelial cell-like cells

The WTC11 human iPSC line (Coriell Institute, GM25256), derived from a healthy male donor, was maintained on growth factor-reduced Matrigel (Corning) in mTeSR Plus medium (STEMCELL Technologies) supplemented with 1% penicillin-streptomycin (Nacalai Tesque) at 37°C in a humidified incubator with 5% CO₂. The medium was changed every other day, and cells were passaged at approximately 80% confluence. For passaging, cells were dissociated with Accutase (Nacalai Tesque) at 37°C for 5 min. For low-density plating, 10 µM Y-27632 (Fujifilm Wako Pure Chemical) was added to the culture medium.

A total of 1.0 × 10⁶ human iPSCs were resuspended in DMEM/F-12 with GlutaMAX (Gibco) supplemented with 2% B-27 without vitamin A (Gibco), 1% N-2 supplement (Gibco), 10 µM SB431542 (Fujifilm Wako Pure Chemical), 0.1 µM LDN193189 (Tocris Bioscience), 10 µM Y-27632, and 1% penicillin-streptomycin, and cultured in suspension in 25-cm² flasks (TPP) for 5 days to induce neurosphere formation.

Neurospheres were then plated onto plates sequentially coated with poly-L-ornithine (Fujifilm Wako Pure Chemical) and laminin (Gibco). When the cultures reached approximately 90% confluence, cells were passaged at a 1:2 ratio and maintained in DMEM/F-12 with GlutaMAX supplemented with 2% B-27 without vitamin A, 1% N-2 supplement, 20 ng/mL FGF2 (PeproTech), 10 µM Y-27632, and 1% penicillin-streptomycin to generate neural progenitor cells (NPCs). After 8 days of culture, neural rosette-like structures were observed.

NPCs (1.45 × 10⁵ cells) were subsequently replated onto Matrigel-coated plates and differentiated into astrocytes by culture in Astrocyte Medium (ScienCell) for 30 days^57^. From day 31 onward, cells were maintained in Astrocyte Medium without astrocyte growth supplement (AGS; ScienCell). For passaging of both NPCs and astrocytes, cells were dissociated with Accumax (Innovative Cell Technologies) at 37°C for 5 min.

For generation of EECM-derived brain microvascular endothelial cell-like cells (EECM-BMEC-like cells), we used two healthy control hiPSC lines, HPS1006 and HPS4290, originally established from healthy donors (36-year-old male and 36-year-old female, respectively) as previously reported^58^. These lines were obtained from the RIKEN BioResource Research Center (RIKEN BRC) through the National BioResource Project of MEXT/AMED, Japan. hiPSCs were cultured on Matrigel-coated plates in mTeSR1 medium (STEMCELL Technologies) under standard humidified conditions at 37°C with 5% CO₂. The culture medium was refreshed daily. Cells were passaged when they reached approximately 80% confluence. For routine passage, colonies were detached using ReLeSR (STEMCELL Technologies) according to the manufacturer’s instructions and transferred onto newly Matrigel-coated plates.

EECM-BMEC-like cells were generated from hiPSCs using the extended endothelial cell culture method (EECM), with minor modifications from our previous reports^59–61^. Briefly, endothelial differentiation was induced by CHIR99021, and CD31-positive cells were isolated by magnetic-activated cell sorting and plated on collagen-coated dishes. Endothelial cells were further purified by repeated brief Accutase treatment based on their preferential detachment from contaminating smooth muscle-like cells. The resulting EECM-BMEC-like cells were used at passages 3-6 and maintained in endothelial maintenance medium (hECSR; human endothelial SFM supplemented with B-27 and 20 ng/mL FGF2) with medium changes every other day.

### AAV transduction assay in iPSC-derived astrocytes and EECM-BMEC-like cells

Human iPSC-derived astrocytes (Fig. 5F, G) were seeded at 1 × 10^4^ cells per well onto 13-mm coverslips (MATSUNAMI) placed in 24-well plates. Coverslips were coated with Matrigel (Corning) at 37°C for 1 h. Cells were seeded at differentiation day 34 and maintained in Astrocyte Medium without astrocyte growth supplement (AGS). Twenty-four hours after seeding (day 35), AAV-AST vectors carrying Region Ref., Region d, or Region dlZ upstream of mCherry were added at 1 × 10^10^ vg per well. Cells were fixed 6 days after transduction and counterstained with Hoechst before fluorescence imaging.

Human iPSC-derived EECM-BMEC-like cells (Fig. 5H, I) were seeded at 2,500 cells per well in collagen-precoated 384-well plates (Corning) and cultured in hECSR medium. Twenty-four hours later, AAV-X1.1 vectors carrying Region Ref., Region d, or Region dlZ upstream of mCherry were added at 5 × 10^9^ vg per well. Live-cell images were acquired 8 days after transduction using a BZ-X810 microscope (KEYENCE, Osaka, Japan) at ×20 magnification. To reduce potential selection bias, images were automatically captured at the center of each well as reported previously^62^.

For fluorescence quantification in the EECM-BMEC-like cell assay, one image was automatically acquired from the center of each well, and 12 wells were analyzed per experiment. mCherry signal was quantified using the BZ-X800 Analyzer software (KEYENCE) as R (integrated), defined as the sum of red-channel pixel intensities within the analyzed image area. Because this value reflects integrated fluorescence intensity rather than an absolute physical quantity, data are presented in arbitrary units (a.u.).

### Statistics

Statistical analyses were performed using GraphPad Prism 10.4.1 (GraphPad Software). Data are presented as mean ± SEM. The statistical tests used are specified in the corresponding figure legends. For comparisons between two groups, two-tailed Welch’s t-tests were used. Normality was assessed using the D’Agostino–Pearson omnibus test or Shapiro–Wilk test, as appropriate. A P value of less than 0.05 was considered statistically significant. Outliers were defined a priori as values exceeding 2 SD from the group mean and were excluded from the analysis. A total of 127 mice were used in this study.

### Study approval

The use of human postmortem brain tissues was approved by the Ethics Committee of Tokyo Metropolitan Institute of Medical Science (no. 23-26), and written informed consent was obtained from the next of kin. All animal procedures were approved by the Animal Experimentation Ethics Committee of the Tokyo Metropolitan Institute of Medical Science (no. 24-034).

## Data availability

Values underlying the graphed data are provided in the Supporting Data Values file. Additional data and analytic code are available from the corresponding author upon reasonable request.

## Acknowledgements

We thank Dr. T. Hara (TMiMS, Tokyo, Japan) for kindly sharing Tie2-Cre mice, originally obtained from The Jackson Laboratory. We are also grateful to Dr. M. Murakami (The University of Tokyo, Tokyo, Japan) for generously providing CAG-Cre mice, originally generated by Dr. J. Miyazaki (Osaka University, Suita, Japan). We further thank Dr. M. Hasegawa (TMiMS, Tokyo, Japan) and Dr. A. Nagakura (Tokyo Metropolitan Matsuzawa Hospital, Tokyo, Japan) for coordinating the acquisition and distribution of postmortem brain samples. We also appreciate Dr. H. Kawaji (TMiMS, Tokyo, Japan) for his assistance in exploring the GLUT1 expression regulatory regions. Finally, we express our deepest gratitude to the families of the deceased individuals for their generous donation of brain tissue and for their time and cooperation during the consent and interview process.

## Author contributions

S.H. designed the study. S.T., H.S., N.A., M.K., Y.Y., A.S., K.M., H.N., E.S., K.S., and S.H. conducted experiments and acquired data. S.T., H.N., and S.H. analyzed data and images. Y.M. and K.S. provided critical input on assays using iPSC-derived astrocytes and on immunostaining of postmortem brain tissue, respectively. M.K. created the schematic illustration shown in Figure S5. S.T. wrote the first draft of the manuscript. S.T., H.S., N.A., and H.N. wrote the Methods section. H.O. and S.H. reviewed and edited the manuscript. S.H. supervised the study.

## Artificial intelligence tools

OpenAI’s ChatGPT (GPT-5.5 Thinking) was used solely for language editing of the manuscript. No data, analyses, or scientific images were generated or modified using AI. All edited text was reviewed and verified by the authors, who take full responsibility for the content of the manuscript.

