## Supplemental data for "Astrocytic and endothelial GLUT1 restoration improves brain glucose homeostasis in GLUT1 deficiency syndrome"

**Supplementary Fig.1**

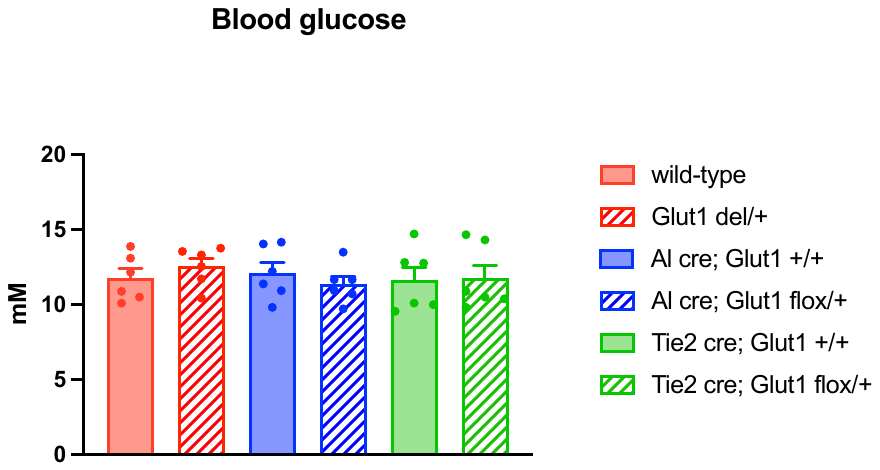
**Supplementary Fig. 1 | Random-fed blood glucose in mice with cell-type–specific Glut1 haploinsufficiency.**

Random-fed (non-fasted) blood glucose concentrations in the indicated genotypes. Bars represent mean ± SEM (n = 6 mice per group). Statistical significance was assessed by one-way ANOVA followed by Tukey’s multiple-comparisons test; no significant differences were observed among groups.

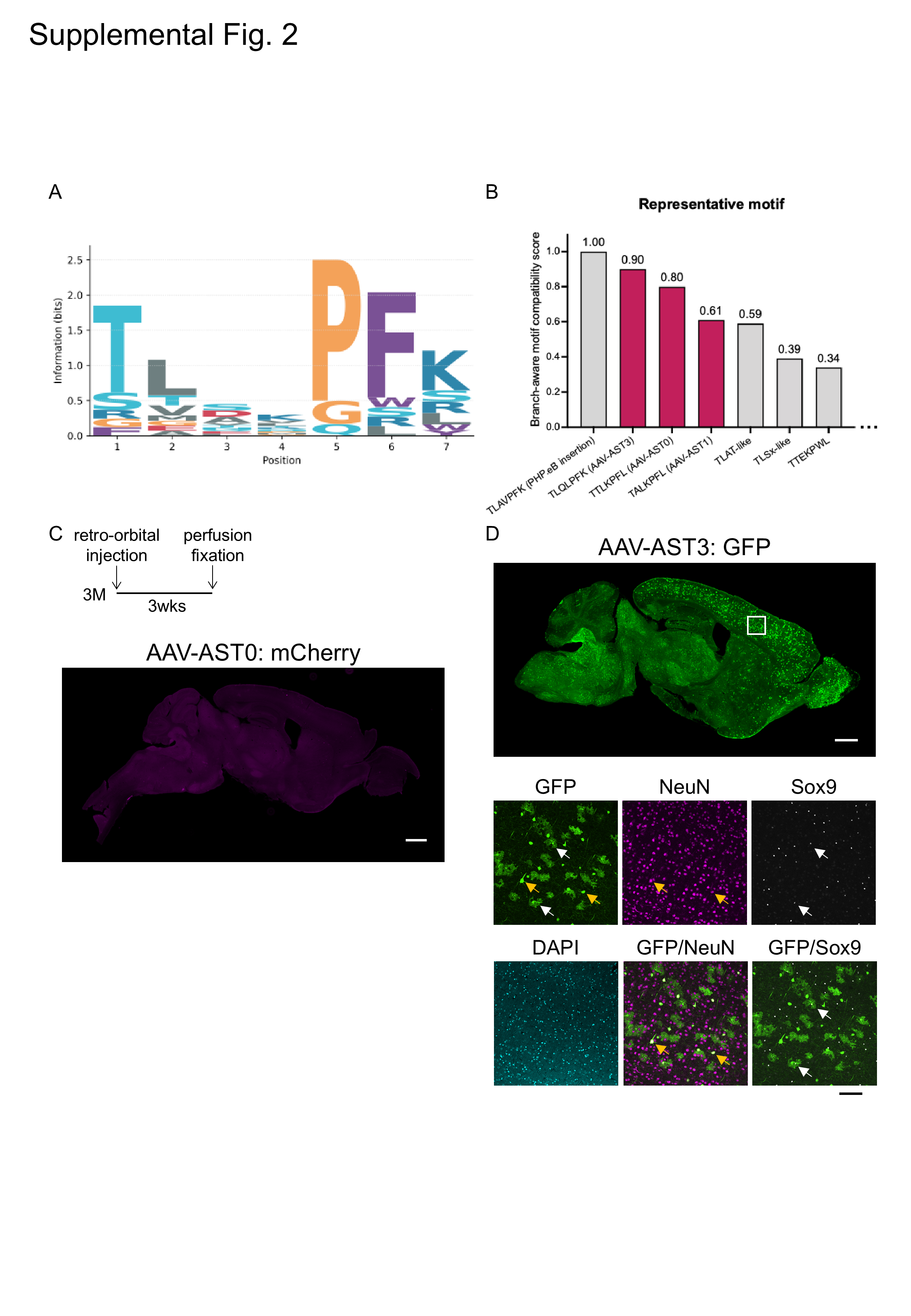
**Supplementary Fig.2**

**Supplementary Fig. 2 | Conserved motif structure and branch-aware ranking of astrocyte/CNS-related AAV insertions.**

(A) Sequence logo representing the positional enrichment pattern of the normalized 7-mer reference motifs used to generate the position-frequency matrix (PFM). Enrichment was observed at Thr (position 1) and Pro-Phe (positions 5-6), delineating a T(L/A/T)xxPF[K/L] core architecture shared by astrocyte/CNS-related variants. (B) Branch-aware motif compatibility scores of representative astrocyte/CNS-related AAV insertion motifs. Four literature-supported PHP.B-like branches (TLAV-like, TALK-like, TTLK-like, and TLQx-like) were defined, and candidate sequences were scored against branch-specific position-frequency models. Scores were normalized to the top-ranked motif (= 1.0). For visualization, near-identical motifs were collapsed and representative sequences are shown. TLAVPFK, TLQLPFK, TTLKPFL, and TALKPFL ranked within the top-scoring group, with TLAVPFK excluded from experimental testing because it is identical to the PHP.eB insertion sequence. (C, D) Representative sagittal brain sections following intravenous administration (retro-orbital injection) of AAV-AST0:CAG-mCherry (C) and AAV-AST3:CAG-GFP (D) into 3-month-old mice (100 µL, 3.3 × 10^11^ vg per mouse). Scale bars, 1,000 µm. Enlarged views in (D) show immunostaining for GFP together with NeuN and SOX9. White and yellow arrows indicate representative SOX9-positive astrocytes and NeuN-positive neurons colocalized with GFP signals, respectively. Scale bars, 100 µm.

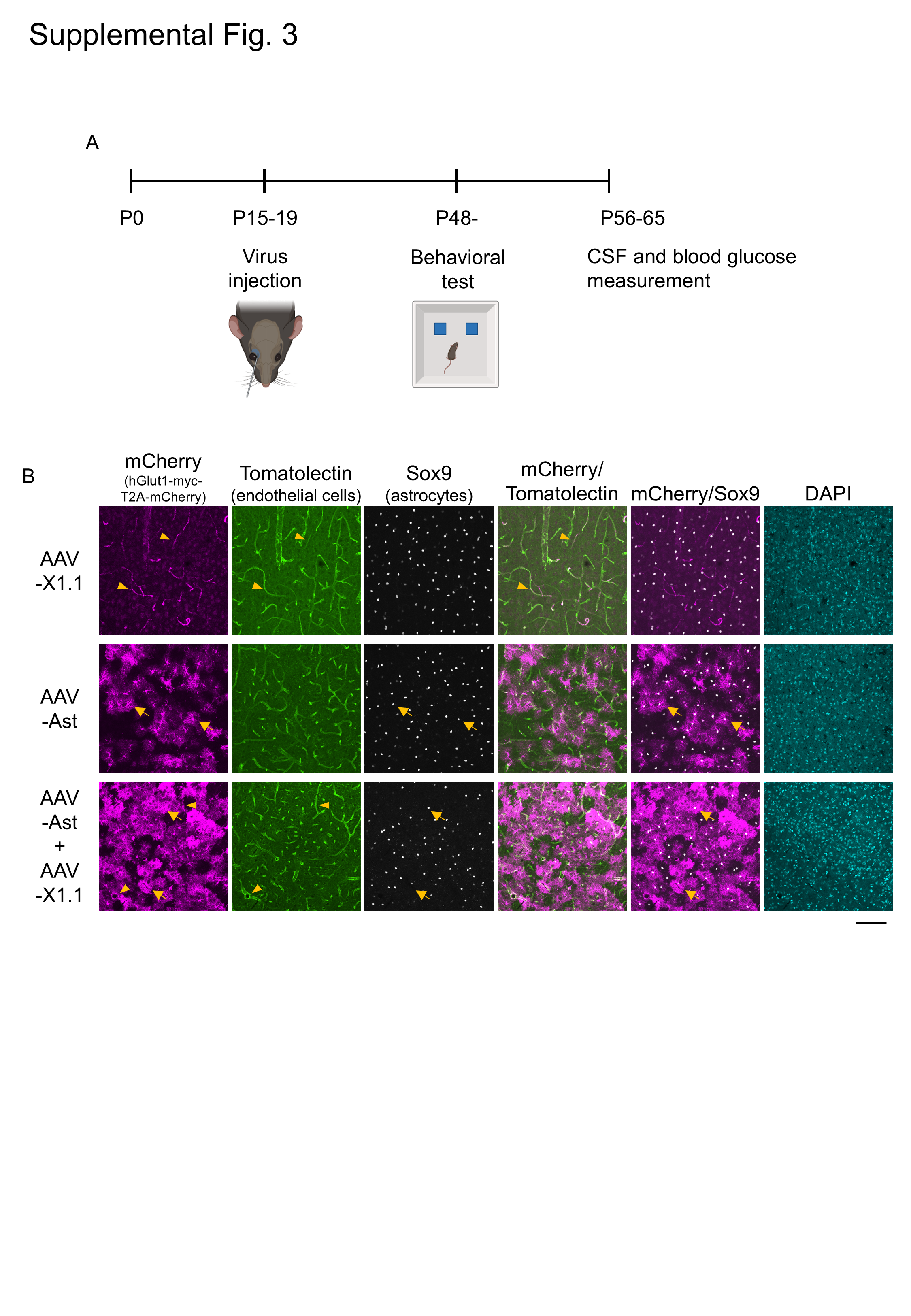
**Supplementary Fig. 3**

**Supplementary Fig. 3 | Cell-type–specific Glut1 supplementation in Glut1 haploinsufficient mice via AAV administration.**

(A) Experimental workflow for Glut1 supplementation using AAV. The illustration was created with BioRender. (B) Representative transgene expression after intravenous delivery of AAV-X1.1 (EC-targeting), AAV-AST (astrocyte-targeting), or AAV-AST + AAV-X1.1 (dual targeting) in Glut1 haploinsufficient mice. Representative confocal z-stack maximum-intensity projections were acquired from the cortex of Glut1 haploinsufficient mice injected with AAV. mCherry is shown in magenta, lectin in green, SOX9 in white, and DAPI in cyan. Yellow arrows and arrowheads indicate representative astrocytes and ECs colocalized with mCherry, respectively. Scale bar, 100 µm.

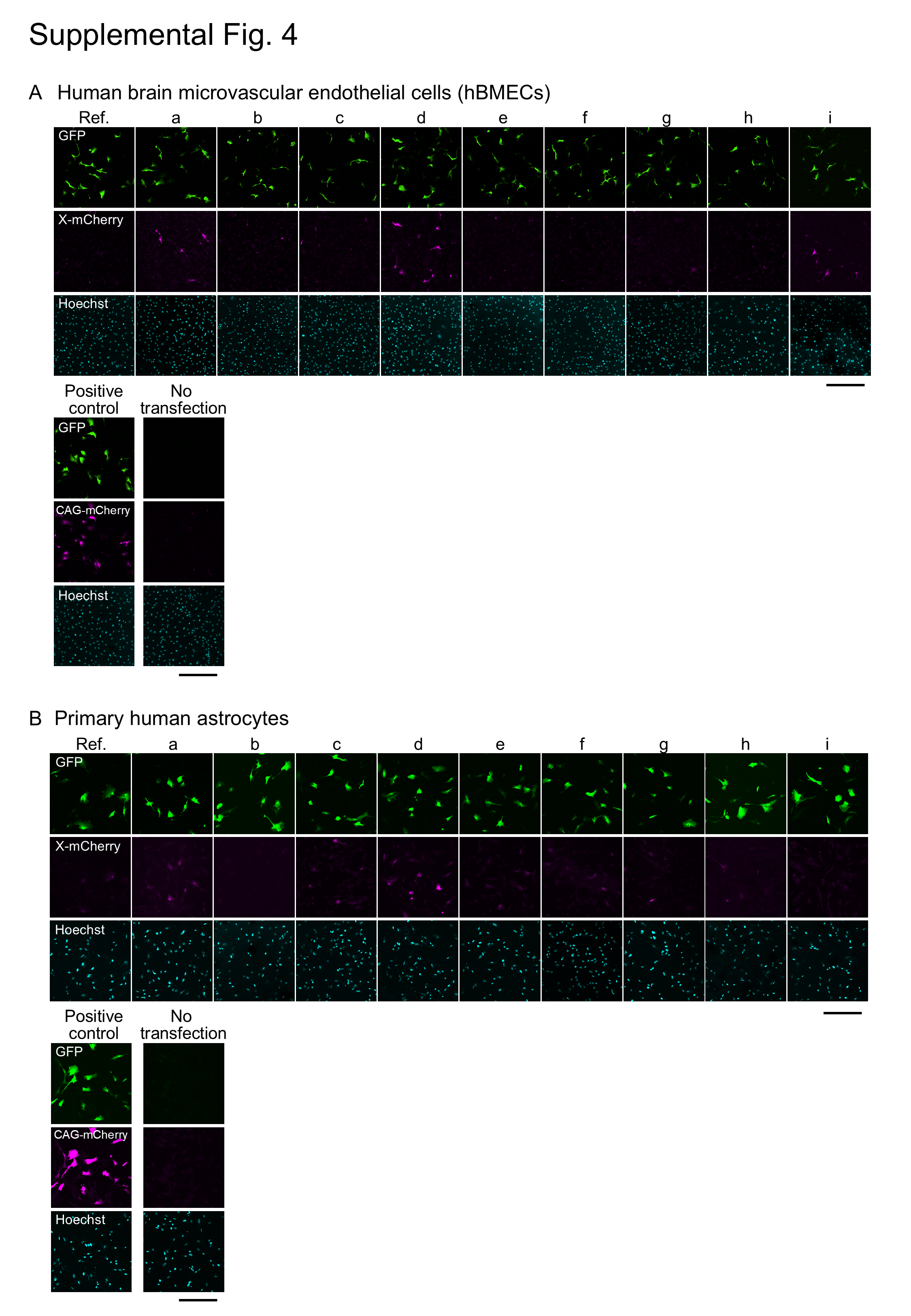
**Supplementary Fig. 4**

**Supplementary Fig. 4 | Functional validation of human SLC2A1 regulatory regions in primary human astrocytes.**

Reporter constructs containing candidate SLC2A1 regulatory regions (a-i) upstream of mCherry were transfected into primary cultured human astrocytes. CAG-GFP was co-transfected to monitor transfection efficiency in all conditions, except for the non-transfected negative control. As a positive control, CAG-mCherry was co-transfected with CAG-GFP. GFP is shown in green, mCherry in magenta, and Hoechst in cyan. Scale bar, 300 µm.

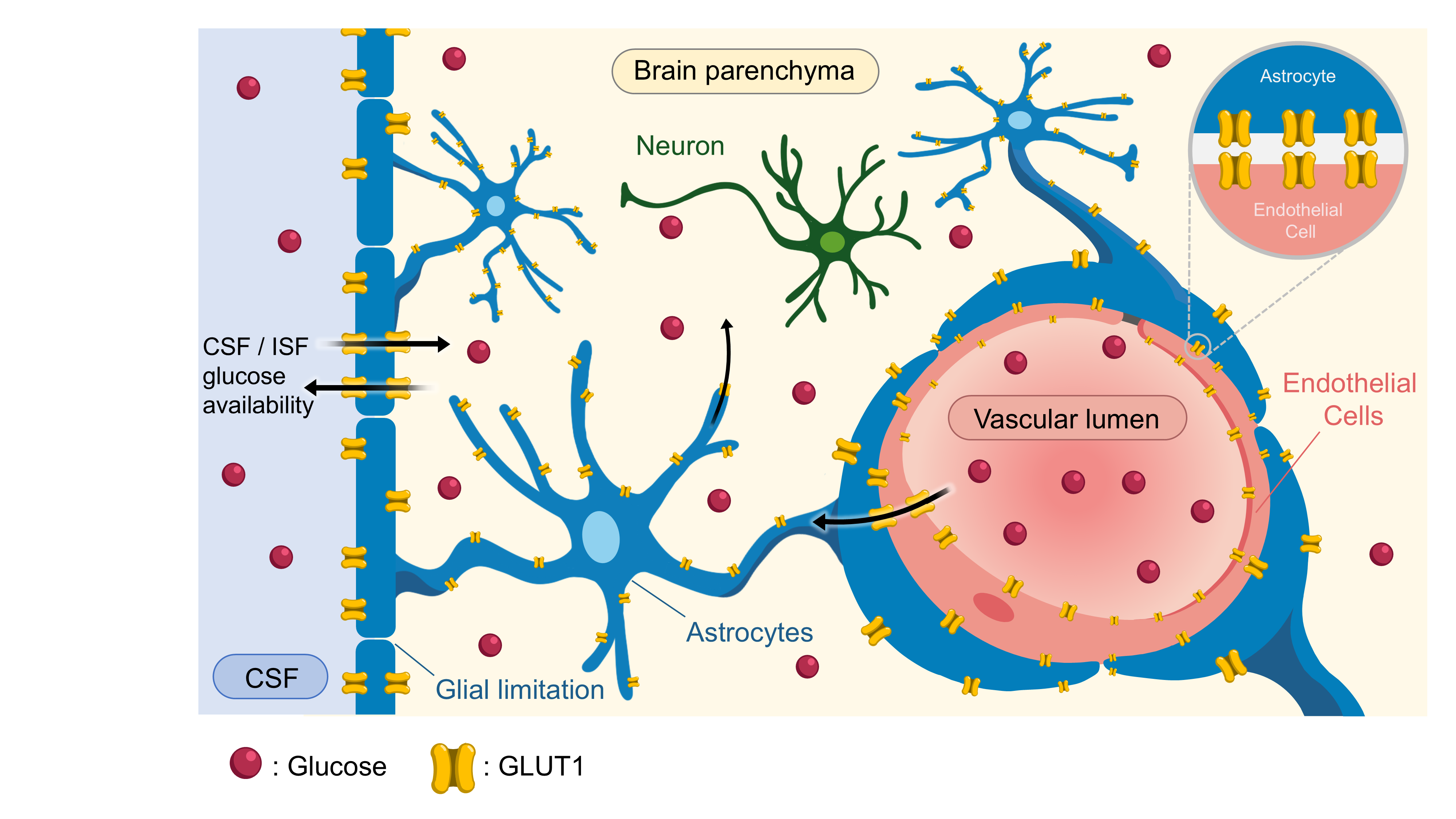
**Supplementary Fig. 5**

**Supplementary Fig. 5 | Model: endothelial and astrocytic GLUT1 form a multicellular gateway for maintaining glucose availability in the brain parenchyma and CSF.**

Endothelial GLUT1 mediates glucose transport across the blood–brain barrier from the vascular lumen into the brain. Astrocytic GLUT1, distributed from perivascular endfeet to astrocytic processes, may further contribute to maintaining glucose availability within the brain parenchyma and, indirectly, the CSF/ISF compartment. Haploinsufficiency of GLUT1 in either endothelial cells or astrocytes is associated with reduced CSF and interstitial fluid glucose levels, as well as cognitive and motor impairments. In our rescue experiments, robust phenotypic improvement required restoration of GLUT1 in both endothelial and astrocytic compartments, supporting a two-compartment model of brain glucose homeostasis. In addition to regulating glucose availability, astrocytes can metabolize glucose into lactate and supply it to neurons via the astrocyte–neuron lactate shuttle. Thus, astrocytes may act as central metabolic regulators that shape intracerebral gradients of two major energy substrates: glucose and lactate.

**Supplementary Table 1. Annotated full nucleotide sequence of pAAV-CAG-GLUT1-myc-T2A-mCherry**

| .**Element** | **Coordinates (nt)** | **Length (bp)** | **Description** |
| --- | --- | --- | --- |
| CAG promoter / regulatory region | 1–1881 | 1881 | CAG promoter region containing CMV enhancer/chicken β-actin-derived regulatory elements |
| XhoI + Kozak | 1882–1893 | 12 | Cloning site and Kozak sequence immediately upstream of the coding region |
| Human GLUT1 (SLC2A1) CDS | 1894–3369 | 1476 | Coding sequence of human GLUT1 |
| myc tag | 3370–3399 | 30 | C-terminal myc epitope tag |
| BamHI | 3400–3405 | 6 | Restriction site |
| T2A | 3406–3462 | 57 | Self-cleaving 2A peptide sequence |
| XhoI + Kozak | 3463–3474 | 12 | Cloning/Kozak sequence upstream of the fluorescent reporter |
| mCherry CDS | 3475–4185 | 711 | Coding sequence of mCherry, including stop codon |
| Plasmid backbone | 4186–6115 | 1930 | Non-expression-cassette backbone sequence |
| bla / AmpR | 6116–6976 | 861 | Beta-lactamase coding sequence for ampicillin resistance |
| Plasmid backbone (ori/noncoding) | 6977–7796 | 820 | Remaining backbone sequence including origin-associated/noncoding region |

**Full nucleotide sequence**

**[1–1881] CAG promoter / regulatory region**

*CAG promoter region containing CMV enhancer/chicken β-actin-derived regulatory elements*

CCTGCAGGCA GCTGCGCGCT CGCTCGCTCA CTGAGGCCGC CCGGGCAAAG CCCGGGCGTC GGGCGACCTT TGGTCGCCCG GCCTCAGTGA GCGAGCGAGC
GCGCAGAGAG GGAGTGGCCA ACTCCATCAC TAGGGGTTCC TTCTAGACTC GACATTGATT ATTGACTAGT TATTAATAGT AATCAATTAC GGGGTCATTA
GTTCATAGCC CATATATGGA GTTCCGCGTT ACATAACTTA CGGTAAATGG CCCGCCTGGC TGACCGCCCA ACGACCCCCG CCCATTGACG TCAATAATGA
CGTATGTTCC CATAGTAACG CCAATAGGGA CTTTCCATTG ACGTCAATGG GTGGAGTATT TACGGTAAAC TGCCCACTTG GCAGTACATC AAGTGTATCA
TATGCCAAGT ACGCCCCCTA TTGACGTCAA TGACGGTAAA TGGCCCGCCT GGCATTATGC CCAGTACATG ACCTTATGGG ACTTTCCTAC TTGGCAGTAC
ATCTACGTAT TAGTCATCGC TATTACCATG GTCGAGGTGA GCCCCACGTT CTGCTTCACT CTCCCCATCT CCCCCCCCTC CCCACCCCCA ATTTTGTATT
TATTTATTTT TTAATTATTT TGTGCAGCGA TGGGGGCGGG GGGGGGGGGG GGGGCGCGCG CCAGGCGGGG CGGGGCGGGG CGAGGGGCGG GGCGGGGCGA
GGCGGAGAGG TGCGGCGGCA GCCAATCAGA GCGGCGCGCT CCGAAAGTTT CCTTTTATGG CGAGGCGGCG GCGGCGGCGG CCCTATAAAA AGCGAAGCGC
GCGGCGGGCG GGAGTCGCTG CGCGCTGCCT TCGCCCCGTG CCCCGCTCCG CCGCCGCCTC GCGCCGCCCG CCCCGGCTCT GACTGACCGC GTTACTCCCA
CAGGTGAGCG GGCGGGACGG CCCTTCTCCT CCGGGCTGTA ATTAGCGCTT GGTTTAATGA CGGCTTGTTT CTTTTCTGTG GCTGCGTGAA AGCCTTGAGG
GGCTCCGGGA GGGCCCTTTG TGCGGGGGGA GCGGCTCGGG GGGTGCGTGC GTGTGTGTGT GCGTGGGGAG CGCCGCGTGC GGCTCCGCGC TGCCCGGCGG
CTGTGAGCGC TGCGGGCGCG GCGCGGGGCT TTGTGCGCTC CGCAGTGTGC GCGAGGGGAG CGCGGCCGGG GGCGGTGCCC CGCGGTGCGG GGGGGGCTGC
GAGGGGAACA AAGGCTGCGT GCGGGGTGTG TGCGTGGGGG GGTGAGCAGG GGGTGTGGGC GCGTCGGTCG GGCTGCAACC CCCCCTGCAC CCCCCTCCCC
GAGTTGCTGA GCACGGCCCG GCTTCGGGTG CGGGGCTCCG TACGGGGCGT GGCGCGGGGC TCGCCGTGCC GGGCGGGGGG TGGCGGCAGG TGGGGGTGCC
GGGCGGGGCG GGGCCGCCTC GGGCCGGGGA GGGCTCGGGG GAGGGGCGCG GCGGCCCCCG GAGCGCCGGC GGCTGTCGAG GCGCGGCGAG CCGCAGCCAT
TGCCTTTTAT GGTAATCGTG CGAGAGGGCG CAGGGACTTC CTTTGTCCCA AATCTGTGCG GAGCCGAAAT CTGGGAGGCG CCGCCGCACC CCCTCTAGCG
GGCGCGGGGC GAAGCGGTGC GGCGCCGGCA GGAAGGAAAT GGGCGGGGAG GGCCTTCGTG CGTCGCCGCG CCGCCGTCCC CTTCTCCCTC TCCAGCCTCG
GGGCTGTCCG CGGGGGGACG GCTGCCTTCG GGGGGGACGG GGCAGGGCGG GGTTCGGCTT CTGGCGTGTG ACCGGCGGCT CTAGAGCCTC TGCTAACCAT
GTTCATGCCT TCTTCTTTTT CCTACAGCTC CTGGGCAACG TGCTGGTTAT TGTGCTGTCT CATCATTTTG GCAAAGAATT G

**[1882–1893] XhoI + Kozak**

*Cloning site and Kozak sequence immediately upstream of the coding region*

CTCGAGGCCA CC

**[1894–3369] Human GLUT1 (SLC2A1) CDS**

*Coding sequence of human GLUT1*

ATGGAGCCCA GCAGCAAGAA GCTGACGGGT CGCCTCATGC TGGCCGTGGG AGGAGCAGTG CTTGGCTCCC TGCAGTTTGG CTACAACACT GGAGTCATCA
ATGCCCCCCA GAAGGTGATC GAGGAGTTCT ACAACCAGAC ATGGGTCCAC CGCTATGGGG AGAGCATCCT GCCCACCACG CTCACCACGC TCTGGTCCCT
CTCAGTGGCC ATCTTTTCTG TTGGGGGCAT GATTGGCTCC TTCTCTGTGG GCCTTTTCGT TAACCGCTTT GGCCGGCGGA ATTCAATGCT GATGATGAAC
CTGCTGGCCT TCGTGTCCGC CGTGCTCATG GGCTTCTCGA AACTGGGCAA GTCCTTTGAG ATGCTGATCC TGGGCCGCTT CATCATCGGT GTGTACTGCG
GCCTGACCAC AGGCTTCGTG CCCATGTATG TGGGTGAAGT GTCACCCACA GCCCTTCGTG GGGCCCTGGG CACCCTGCAC CAGCTGGGCA TCGTCGTCGG
CATCCTCATC GCCCAGGTGT TCGGCCTGGA CTCCATCATG GGCAACAAGG ACCTGTGGCC CCTGCTGCTG AGCATCATCT TCATCCCGGC CCTGCTGCAG
TGCATCGTGC TGCCCTTCTG CCCCGAGAGT CCCCGCTTCC TGCTCATCAA CCGCAACGAG GAGAACCGGG CCAAGAGTGT GCTAAAGAAG CTGCGCGGGA
CAGCTGACGT GACCCATGAC CTGCAGGAGA TGAAGGAAGA GAGTCGGCAG ATGATGCGGG AGAAGAAGGT CACCATCCTG GAGCTGTTCC GCTCCCCCGC
CTACCGCCAG CCCATCCTCA TCGCTGTGGT GCTGCAGCTG TCCCAGCAGC TGTCTGGCAT CAACGCTGTC TTCTATTACT CCACGAGCAT CTTCGAGAAG
GCGGGGGTGC AGCAGCCTGT GTATGCCACC ATTGGCTCCG GTATCGTCAA CACGGCCTTC ACTGTCGTGT CGCTGTTTGT GGTGGAGCGA GCAGGCCGGC
GGACCCTGCA CCTCATAGGC CTCGCTGGCA TGGCGGGTTG TGCCATACTC ATGACCATCG CGCTAGCACT GCTGGAGCAG CTACCCTGGA TGTCCTATCT
GAGCATCGTG GCCATCTTTG GCTTTGTGGC CTTCTTTGAA GTGGGTCCTG GCCCCATCCC ATGGTTCATC GTGGCTGAAC TCTTCAGCCA GGGTCCACGT
CCAGCTGCCA TTGCCGTTGC AGGCTTCTCC AACTGGACCT CAAATTTCAT TGTGGGCATG TGCTTCCAGT ATGTGGAGCA ACTGTGTGGT CCCTACGTCT
TCATCATCTT CACTGTGCTC CTGGTTCTGT TCTTCATCTT CACCTACTTC AAAGTTCCTG AGACTAAAGG CCGGACCTTC GATGAGATCG CTTCCGGCTT
CCGGCAGGGG GGAGCCAGCC AAAGTGACAA GACACCCGAG GAGCTGTTCC ATCCCCTGGG GGCTGATTCC CAAGTG

**[3370–3399] myc tag**

*C-terminal myc epitope tag*

GAACAAAAAC TCATCTCAGA AGAGGATCTG

**[3400–3405] BamHI**

*Restriction site*

GGATCC

**[3406–3462] T2A**

*Self-cleaving 2A peptide sequence*

GGAGAGGGCA GAGGAAGTCT GCTAACATGC GGTGACGTCG AGGAGAATCC TGGCCCA

**[3463–3474] XhoI + Kozak**

*Cloning/Kozak sequence upstream of the fluorescent reporter*

CTCGAGGCCA CC

**[3475–4185] mCherry CDS**

*Coding sequence of mCherry, including stop codon*

ATGGTGAGCA AGGGCGAGGA GGATAACATG GCCATCATCA AGGAGTTCAT GCGCTTCAAG GTGCACATGG AGGGCTCCGT GAACGGCCAC GAGTTCGAGA
TCGAGGGCGA GGGCGAGGGC CGCCCCTACG AGGGCACCCA GACCGCCAAG CTGAAGGTGA CCAAGGGTGG CCCCCTGCCC TTCGCCTGGG ACATCCTGTC
CCCTCAGTTC ATGTACGGCT CCAAGGCCTA CGTGAAGCAC CCCGCCGACA TCCCCGACTA CTTGAAGCTG TCCTTCCCCG AGGGCTTCAA GTGGGAGCGC
GTGATGAACT TCGAGGACGG CGGCGTGGTG ACCGTGACCC AGGACTCCTC CCTGCAGGAC GGCGAGTTCA TCTACAAGGT GAAGCTGCGC GGCACCAACT
TCCCCTCCGA CGGCCCCGTA ATGCAGAAGA AGACCATGGG CTGGGAGGCC TCCTCCGAGC GGATGTACCC CGAGGACGGC GCCCTGAAGG GCGAGATCAA
GCAGAGGCTG AAGCTGAAGG ACGGCGGCCA CTACGACGCT GAGGTCAAGA CCACCTACAA GGCCAAGAAG CCCGTGCAGC TGCCCGGCGC CTACAACGTC
AACATCAAGT TGGACATCAC CTCCCACAAC GAGGACTACA CCATCGTGGA ACAGTACGAA CGCGCCGAGG GCCGCCACTC CACCGGCGGC ATGGACGAGC
TGTACAAGTA A

**[4186–6115] Plasmid backbone**

*Non-expression-cassette backbone sequence*

ACCCAGCTTT CTTGTACAAA GTGGGAATTC CGATAATCAA CCTCTGGATT ACAAAATTTG TGAAAGATTG ACTGGTATTC TTAACTATGT TGCTCCTTTT
ACGCTATGTG GATACGCTGC TTTAATGCCT TTGTATCATG CTATTGCTTC CCGTATGGCT TTCATTTTCT CCTCCTTGTA TAAATCCTGG TTGCTGTCTC
TTTATGAGGA GTTGTGGCCC GTTGTCAGGC AACGTGGCGT GGTGTGCACT GTGTTTGCTG ACGCAACCCC CACTGGTTGG GGCATTGCCA CCACCTGTCA
GCTCCTTTCC GGGACTTTCG CTTTCCCCCT CCCTATTGCC ACGGCGGAAC TCATCGCCGC CTGCCTTGCC CGCTGCTGGA CAGGGGCTCG GCTGTTGGGC
ACTGACAATT CCGTGGTGTT GTCGGGGAAG CTGACGTCCT TTCCATGGCT GCTCGCCTGT GTTGCCACCT GGATTCTGCG CGGGACGTCC TTCTGCTACG
TCCCTTCGGC CCTCAATCCA GCGGACCTTC CTTCCCGCGG CCTGCTGCCG GCTCTGCGGC CTCTTCCGCG TCTTCGCCTT CGCCCTCAGA CGAGTCGGAT
CTCCCTTTGG GCCGCCTCCC CGCATCGGGA ATTCCTAGAG CTCGCTGATC AGCCTCGACT GTGCCTTCTA GTTGCCAGCC ATCTGTTGTT TGCCCCTCCC
CCGTGCCTTC CTTGACCCTG GAAGGTGCCA CTCCCACTGT CCTTTCCTAA TAAAATGAGG AAATTGCATC GCATTGTCTG AGTAGGTGTC ATTCTATTCT
GGGGGGTGGG GTGGGGCAGG ACAGCAAGGG GGAGGATTGG GAAGAGAATA GCAGGCATGC TGGGGAGGGC CGCAGGAACC CCTAGTGATG GAGTTGGCCA
CTCCCTCTCT GCGCGCTCGC TCGCTCACTG AGGCCGGGCG ACCAAAGGTC GCCCGACGCC CGGGCTTTGC CCGGGCGGCC TCAGTGAGCG AGCGAGCGCG
CAGCTGCCTG CAGGGGCGCC TGATGCGGTA TTTTCTCCTT ACGCATCTGT GCGGTATTTC ACACCGCATA CGTCAAAGCA ACCATAGTAC GCGCCCTGTA
GCGGCGCATT AAGCGCGGCG GGGGTGGTGG TTACGCGCAG CGTGACCGCT ACACTTGCCA GCGCCTTAGC GCCCGCTCCT TTCGCTTTCT TCCCTTCCTT
TCTCGCCACG TTCGCCGGCT TTCCCCGTCA AGCTCTAAAT CGGGGGCTCC CTTTAGGGTT CCGATTTAGT GCTTTACGGC ACCTCGACCC CAAAAAACTT
GATTTGGGTG ATGGTTCACG TAGTGGGCCA TCGCCCTGAT AGACGGTTTT TCGCCCTTTG ACGTTGGAGT CCACGTTCTT TAATAGTGGA CTCTTGTTCC
AAACTGGAAC AACACTCAAC TCTATCTCGG GCTATTCTTT TGATTTATAA GGGATTTTGC CGATTTCGGT CTATTGGTTA AAAAATGAGC TGATTTAACA
AAAATTTAAC GCGAATTTTA ACAAAATATT AACGTTTACA ATTTTATGGT GCACTCTCAG TACAATCTGC TCTGATGCCG CATAGTTAAG CCAGCCCCGA
CACCCGCCAA CACCCGCTGA CGCGCCCTGA CGGGCTTGTC TGCTCCCGGC ATCCGCTTAC AGACAAGCTG TGACCGTCTC CGGGAGCTGC ATGTGTCAGA
GGTTTTCACC GTCATCACCG AAACGCGCGA GACGAAAGGG CCTCGTGATA CGCCTATTTT TATAGGTTAA TGTCATGATA ATAATGGTTT CTTAGACGTC
AGGTGGCACT TTTCGGGGAA ATGTGCGCGG AACCCCTATT TGTTTATTTT TCTAAATACA TTCAAATATG TATCCGCTCA TGAGACAATA ACCCTGATAA
ATGCTTCAAT AATATTGAAA AAGGAAGAGT

**[6116–6976] bla / AmpR**

*Beta-lactamase coding sequence for ampicillin resistance*

ATGAGTATTC AACATTTCCG TGTCGCCCTT ATTCCCTTTT TTGCGGCATT TTGCCTTCCT GTTTTTGCTC ACCCAGAAAC GCTGGTGAAA GTAAAAGATG
CTGAAGATCA GTTGGGTGCA CGAGTGGGTT ACATCGAACT GGATCTCAAC AGCGGTAAGA TCCTTGAGAG TTTTCGCCCC GAAGAACGTT TTCCAATGAT
GAGCACTTTT AAAGTTCTGC TATGTGGCGC GGTATTATCC CGTATTGACG CCGGGCAAGA GCAACTCGGT CGCCGCATAC ACTATTCTCA GAATGACTTG
GTTGAGTACT CACCAGTCAC AGAAAAGCAT CTTACGGATG GCATGACAGT AAGAGAATTA TGCAGTGCTG CCATAACCAT GAGTGATAAC ACTGCGGCCA
ACTTACTTCT GACAACGATC GGAGGACCGA AGGAGCTAAC CGCTTTTTTG CACAACATGG GGGATCATGT AACTCGCCTT GATCGTTGGG AACCGGAGCT
GAATGAAGCC ATACCAAACG ACGAGCGTGA CACCACGATG CCTGTAGCAA TGGCAACAAC GTTGCGCAAA CTATTAACTG GCGAACTACT TACTCTAGCT
TCCCGGCAAC AATTAATAGA CTGGATGGAG GCGGATAAAG TTGCAGGACC ACTTCTGCGC TCGGCCCTTC CGGCTGGCTG GTTTATTGCT GATAAATCTG
GAGCCGGTGA GCGTGGAAGC CGCGGTATCA TTGCAGCACT GGGGCCAGAT GGTAAGCCCT CCCGTATCGT AGTTATCTAC ACGACGGGGA GTCAGGCAAC
TATGGATGAA CGAAATAGAC AGATCGCTGA GATAGGTGCC TCACTGATTA AGCATTGGTA A

**[6977–7796] Plasmid backbone (ori/noncoding)**

*Remaining backbone sequence including origin-associated/noncoding region*

CTGTCAGACC AAGTTTACTC ATATATACTT TAGATTGATT TAAAACTTCA TTTTTAATTT AAAAGGATCT AGGTGAAGAT CCTTTTTGAT AATCTCATGA
CCAAAATCCC TTAACGTGAG TTTTCGTTCC ACTGAGCGTC AGACCCCGTA GAAAAGATCA AAGGATCTTC TTGAGATCCT TTTTTTCTGC GCGTAATCTG
CTGCTTGCAA ACAAAAAAAC CACCGCTACC AGCGGTGGTT TGTTTGCCGG ATCAAGAGCT ACCAACTCTT TTTCCGAAGG TAACTGGCTT CAGCAGAGCG
CAGATACCAA ATACTGTTCT TCTAGTGTAG CCGTAGTTAG GCCACCACTT CAAGAACTCT GTAGCACCGC CTACATACCT CGCTCTGCTA ATCCTGTTAC
CAGTGGCTGC TGCCAGTGGC GATAAGTCGT GTCTTACCGG GTTGGACTCA AGACGATAGT TACCGGATAA GGCGCAGCGG TCGGGCTGAA CGGGGGGTTC
GTGCACACAG CCCAGCTTGG AGCGAACGAC CTACACCGAA CTGAGATACC TACAGCGTGA GCTATGAGAA AGCGCCACGC TTCCCGAAGG GAGAAAGGCG
GACAGGTATC CGGTAAGCGG CAGGGTCGGA ACAGGAGAGC GCACGAGGGA GCTTCCAGGG GGAAACGCCT GGTATCTTTA TAGTCCTGTC GGGTTTCGCC
ACCTCTGACT TGAGCGTCGA TTTTTGTGAT GCTCGTCAGG GGGGCGGAGC CTATGGAAAA ACGCCAGCAA CGCGGCCTTT TTACGGTTCC TGGCCTTTTG
CTGGCCTTTT GCTCACATGT

**Supplementary Table 2. Annotated full nucleotide sequence of pAAV-CAG-mCherry-spacer-WPRE**

| **Element** | **Coordinates (nt)** | **Length (bp)** | **Description** |
| --- | --- | --- | --- |
| 5′ AAV2 ITR | 1–145 | 145 | 5′ inverted terminal repeat |
| CAG promoter / regulatory region | 146–1881 | 1736 | CAG promoter region containing CMV enhancer/chicken β-actin-derived regulatory elements |
| XbaI (ITR/CAG junction) | 142–147 | 6 | Restriction site overlapping the 5′ ITR/CAG junction |
| XbaI | 1779–1784 | 6 | Restriction site in the downstream promoter-proximal region |
| XhoI + Kozak | 1882–1893 | 12 | Cloning site and Kozak sequence immediately upstream of mCherry |
| mCherry CDS | 1894–2598 | 705 | Coding sequence of mCherry, including stop codon |
| Spacer | 2599–3373 | 775 | Inserted spacer sequence between mCherry and WPRE |
| WPRE | 3374–3967 | 594 | Woodchuck hepatitis virus posttranscriptional regulatory element |
| Intervening noncoding region | 3968–4208 | 241 | Noncoding sequence between WPRE and the 3′ ITR |
| 3′ AAV2 ITR | 4209–4353 | 145 | 3′ inverted terminal repeat |
| Plasmid backbone | 4354–5269 | 916 | Non-expression-cassette backbone sequence |
| bla / AmpR | 5270–6130 | 861 | Beta-lactamase coding sequence for ampicillin resistance |
| Plasmid backbone (ori/noncoding) | 6131–6950 | 820 | Remaining backbone sequence including origin-associated/noncoding region |

**Full nucleotide sequence**

**[1–145] 5′ AAV2 ITR**

*5′ inverted terminal repeat*

CCTGCAGGCA GCTGCGCGCT CGCTCGCTCA CTGAGGCCGC CCGGGCAAAG CCCGGGCGTC GGGCGACCTT TGGTCGCCCG
GCCTCAGTGA GCGAGCGAGC GCGCAGAGAG GGAGTGGCCA ACTCCATCAC TAGGGGTTCC TTCTA

**[146–1881] CAG promoter / regulatory region**

*CAG promoter region containing CMV enhancer/chicken β-actin-derived regulatory elements*

GACTCGACAT TGATTATTGA CTAGTTATTA ATAGTAATCA ATTACGGGGT CATTAGTTCA TAGCCCATAT ATGGAGTTCC
GCGTTACATA ACTTACGGTA AATGGCCCGC CTGGCTGACC GCCCAACGAC CCCCGCCCAT TGACGTCAAT AATGACGTAT
GTTCCCATAG TAACGCCAAT AGGGACTTTC CATTGACGTC AATGGGTGGA GTATTTACGG TAAACTGCCC ACTTGGCAGT
ACATCAAGTG TATCATATGC CAAGTACGCC CCCTATTGAC GTCAATGACG GTAAATGGCC CGCCTGGCAT TATGCCCAGT
ACATGACCTT ATGGGACTTT CCTACTTGGC AGTACATCTA CGTATTAGTC ATCGCTATTA CCATGGTCGA GGTGAGCCCC
ACGTTCTGCT TCACTCTCCC CATCTCCCCC CCCTCCCCAC CCCCAATTTT GTATTTATTT ATTTTTTAAT TATTTTGTGC
AGCGATGGGG GCGGGGGGGG GGGGGGGGCG CGCGCCAGGC GGGGCGGGGC GGGGCGAGGG GCGGGGCGGG GCGAGGCGGA
GAGGTGCGGC GGCAGCCAAT CAGAGCGGCG CGCTCCGAAA GTTTCCTTTT ATGGCGAGGC GGCGGCGGCG GCGGCCCTAT
AAAAAGCGAA GCGCGCGGCG GGCGGGAGTC GCTGCGCGCT GCCTTCGCCC CGTGCCCCGC TCCGCCGCCG CCTCGCGCCG
CCCGCCCCGG CTCTGACTGA CCGCGTTACT CCCACAGGTG AGCGGGCGGG ACGGCCCTTC TCCTCCGGGC TGTAATTAGC
GCTTGGTTTA ATGACGGCTT GTTTCTTTTC TGTGGCTGCG TGAAAGCCTT GAGGGGCTCC GGGAGGGCCC TTTGTGCGGG
GGGAGCGGCT CGGGGGGTGC GTGCGTGTGT GTGTGCGTGG GGAGCGCCGC GTGCGGCTCC GCGCTGCCCG GCGGCTGTGA
GCGCTGCGGG CGCGGCGCGG GGCTTTGTGC GCTCCGCAGT GTGCGCGAGG GGAGCGCGGC CGGGGGCGGT GCCCCGCGGT
GCGGGGGGGG CTGCGAGGGG AACAAAGGCT GCGTGCGGGG TGTGTGCGTG GGGGGGTGAG CAGGGGGTGT GGGCGCGTCG
GTCGGGCTGC AACCCCCCCT GCACCCCCCT CCCCGAGTTG CTGAGCACGG CCCGGCTTCG GGTGCGGGGC TCCGTACGGG
GCGTGGCGCG GGGCTCGCCG TGCCGGGCGG GGGGTGGCGG CAGGTGGGGG TGCCGGGCGG GGCGGGGCCG CCTCGGGCCG
GGGAGGGCTC GGGGGAGGGG CGCGGCGGCC CCCGGAGCGC CGGCGGCTGT CGAGGCGCGG CGAGCCGCAG CCATTGCCTT
TTATGGTAAT CGTGCGAGAG GGCGCAGGGA CTTCCTTTGT CCCAAATCTG TGCGGAGCCG AAATCTGGGA GGCGCCGCCG
CACCCCCTCT AGCGGGCGCG GGGCGAAGCG GTGCGGCGCC GGCAGGAAGG AAATGGGCGG GGAGGGCCTT CGTGCGTCGC
CGCGCCGCCG TCCCCTTCTC CCTCTCCAGC CTCGGGGCTG TCCGCGGGGG GACGGCTGCC TTCGGGGGGG ACGGGGCAGG
GCGGGGTTCG GCTTCTGGCG TGTGACCGGC GGCTCTAGAG CCTCTGCTAA CCATGTTCAT GCCTTCTTCT TTTTCCTACA
GCTCCTGGGC AACGTGCTGG TTATTGTGCT GTCTCATCAT TTTGGCAAAG AATTGG

**[142–147] XbaI (ITR/CAG junction)**

*Restriction site overlapping the 5′ ITR/CAG junction*

TCTAGA

**[1779–1784] XbaI**

*Restriction site in the downstream promoter-proximal region*

TCTAGA

**[1882–1893] XhoI + Kozak**

*Cloning site and Kozak sequence immediately upstream of mCherry*

CTCGAGGCCA CC

**[1894–2598] mCherry CDS**

*Coding sequence of mCherry, including stop codon*

ATGGTGAGCA AGGGCGAGGA GGTCATCAAA GAGTTCATGC GCTTCAAGGT GCGCATGGAG GGCTCCATGA ACGGCCACGA
GTTCGAGATC GAGGGCGAGG GCGAGGGCCG CCCCTACGAG GGCACCCAGA CCGCCAAGCT GAAGGTGACC AAGGGCGGCC
CCCTGCCCTT CGCCTGGGAC ATCCTGTCCC CCCAGTTCAT GTACGGCTCC AAGGCGTACG TGAAGCACCC CGCCGACATC
CCCGATTACA AGAAGCTGTC CTTCCCCGAG GGCTTCAAGT GGGAGCGCGT GATGAACTTC GAGGACGGCG GTCTGGTGAC
CGTGACCCAG GACTCCTCCC TGCAGGACGG CACGCTGATC TACAAGGTGA AGATGCGCGG CACCAACTTC CCCCCCGACG
GCCCCGTAAT GCAGAAGAAG ACCATGGGCT GGGAGGCCTC CACCGAGCGC CTGTACCCCC GCGACGGCGT GCTGAAGGGC
GAGATCCACC AGGCCCTGAA GCTGAAGGAC GGCGGCCACT ACCTGGTGGA GTTCAAGACC ATCTACATGG CCAAGAAGCC
CGTGCAACTG CCCGGCTACT ACTACGTGGA CACCAAGCTG GACATCACCT CCCACAACGA GGACTACACC ATCGTGGAAC
AGTACGAGCG CTCCGAGGGC CGCCACCACC TGTTCCTGTA CGGCATGGAC GAGCTGTACA AGTGA

**[2599–3373] Spacer**

*Inserted spacer sequence between mCherry and WPRE*

ACCCAGCTTT CTTGTACAAA GTGGGCATTA CTTATTGTTT TAGCTGTCCT CATGAATGTC TTTTCACTAC CCATTTGCTT
ATCCTGCATC TCTCAGCCTT GACTCCACTC AGTTCTCTTG CTTAGAGATA CCACCTTTCC CCTGAAGTGT TCCTTCCATG
TTTTACGGCG AGATGGTTTC TCCTCGCCTG GGTCGAGCAG TGTGGTTTTG CAAGAGGAAG CAAAAAGCCT CTCCACCCAG
GCCTGGAATG TTTCCACCCA ATGTCGAGCA GTGTGGTTTT GCAAGAGGAA GCAAAAAGCC TCTCCACCCA GGCCTGGAAT
GTTTCCACCC AATGTCGAGC AAACCCCGCC CAGCGTCTTG TCATTGGCGA ATTCGAACAC GCAGATGCAG TCGGGGCGGC
GCGGTCCCAG GTCCACTTCG CATATTAAGG TGACGCGTGT GGCCTCGAAC ACCGAGCGAC CCTGCAGCCA ATATGGGATC
GGCCATTGAA CAAGATGGAT TGCACGCAGG TTCTCCGGCC GCTTGGGTGG AGAGGCTATT CGGCTATGAC TGGGCACAAC
AGACAATCGG CTGCTCTGAT GCCGCCGTGT TCCGGCTGTC AGCGCAGGGG CGCCCGGTTC TTTTTGTCAA GACCGACCTG
TCCGGTGCCC TGAATGAACT GCAGGACGAG GCAGCGCGGC TATCGTGGCT GGCCACGACG GGCGTTCCTT GCGCAGCTGT
GCTCGACGTT GTCACTGAGC TCGAATTCGA TATCAAGCTT ATCGATAATT CCGAT

**[3374–3967] WPRE**

*Woodchuck hepatitis virus posttranscriptional regulatory element*

AATCAACCTC TGGATTACAA AATTTGTGAA AGATTGACTG GTATTCTTAA CTATGTTGCT CCTTTTACGC TATGTGGATA
CGCTGCTTTA ATGCCTTTGT ATCATGCTAT TGCTTCCCGT ATGGCTTTCA TTTTCTCCTC CTTGTATAAA TCCTGGTTGC
TGTCTCTTTA TGAGGAGTTG TGGCCCGTTG TCAGGCAACG TGGCGTGGTG TGCACTGTGT TTGCTGACGC AACCCCCACT
GGTTGGGGCA TTGCCACCAC CTGTCAGCTC CTTTCCGGGA CTTTCGCTTT CCCCCTCCCT ATTGCCACGG CGGAACTCAT
CGCCGCCTGC CTTGCCCGCT GCTGGACAGG GGCTCGGCTG TTGGGCACTG ACAATTCCGT GGTGTTGTCG GGGAAGCTGA
CGTCCTTTCC ATGGCTGCTC GCCTGTGTTG CCACCTGGAT TCTGCGCGGG ACGTCCTTCT GCTACGTCCC TTCGGCCCTC
AATCCAGCGG ACCTTCCTTC CCGCGGCCTG CTGCCGGCTC TGCGGCCTCT TCCGCGTCTT CGCCTTCGCC CTCAGACGAG
TCGGATCTCC CTTTGGGCCG CCTCCCCGCA TCGG

**[3968–4208] Intervening noncoding region**

*Noncoding sequence between WPRE and the 3′ ITR*

GAATTCCTAG AGCTCGCTGA TCAGCCTCGA CTGTGCCTTC TAGTTGCCAG CCATCTGTTG TTTGCCCCTC CCCCGTGCCT
TCCTTGACCC TGGAAGGTGC CACTCCCACT GTCCTTTCCT AATAAAATGA GGAAATTGCA TCGCATTGTC TGAGTAGGTG
TCATTCTATT CTGGGGGGTG GGGTGGGGCA GGACAGCAAG GGGGAGGATT GGGAAGAGAA TAGCAGGCAT GCTGGGGAGG
G

**[4209–4353] 3′ AAV2 ITR**

*3′ inverted terminal repeat*

CCGCAGGAAC CCCTAGTGAT GGAGTTGGCC ACTCCCTCTC TGCGCGCTCG CTCGCTCACT GAGGCCGGGC GACCAAAGGT
CGCCCGACGC CCGGGCTTTG CCCGGGCGGC CTCAGTGAGC GAGCGAGCGC GCAGCTGCCT GCAGG

**[4354–5269] Plasmid backbone**

*Non-expression-cassette backbone sequence*

GGCGCCTGAT GCGGTATTTT CTCCTTACGC ATCTGTGCGG TATTTCACAC CGCATACGTC AAAGCAACCA TAGTACGCGC
CCTGTAGCGG CGCATTAAGC GCGGCGGGGG TGGTGGTTAC GCGCAGCGTG ACCGCTACAC TTGCCAGCGC CTTAGCGCCC
GCTCCTTTCG CTTTCTTCCC TTCCTTTCTC GCCACGTTCG CCGGCTTTCC CCGTCAAGCT CTAAATCGGG GGCTCCCTTT
AGGGTTCCGA TTTAGTGCTT TACGGCACCT CGACCCCAAA AAACTTGATT TGGGTGATGG TTCACGTAGT GGGCCATCGC
CCTGATAGAC GGTTTTTCGC CCTTTGACGT TGGAGTCCAC GTTCTTTAAT AGTGGACTCT TGTTCCAAAC TGGAACAACA
CTCAACTCTA TCTCGGGCTA TTCTTTTGAT TTATAAGGGA TTTTGCCGAT TTCGGTCTAT TGGTTAAAAA ATGAGCTGAT
TTAACAAAAA TTTAACGCGA ATTTTAACAA AATATTAACG TTTACAATTT TATGGTGCAC TCTCAGTACA ATCTGCTCTG
ATGCCGCATA GTTAAGCCAG CCCCGACACC CGCCAACACC CGCTGACGCG CCCTGACGGG CTTGTCTGCT CCCGGCATCC
GCTTACAGAC AAGCTGTGAC CGTCTCCGGG AGCTGCATGT GTCAGAGGTT TTCACCGTCA TCACCGAAAC GCGCGAGACG
AAAGGGCCTC GTGATACGCC TATTTTTATA GGTTAATGTC ATGATAATAA TGGTTTCTTA GACGTCAGGT GGCACTTTTC
GGGGAAATGT GCGCGGAACC CCTATTTGTT TATTTTTCTA AATACATTCA AATATGTATC CGCTCATGAG ACAATAACCC
TGATAAATGC TTCAATAATA TTGAAAAAGG AAGAGT

**[5270–6130] bla / AmpR**

*Beta-lactamase coding sequence for ampicillin resistance*

ATGAGTATTC AACATTTCCG TGTCGCCCTT ATTCCCTTTT TTGCGGCATT TTGCCTTCCT GTTTTTGCTC ACCCAGAAAC
GCTGGTGAAA GTAAAAGATG CTGAAGATCA GTTGGGTGCA CGAGTGGGTT ACATCGAACT GGATCTCAAC AGCGGTAAGA
TCCTTGAGAG TTTTCGCCCC GAAGAACGTT TTCCAATGAT GAGCACTTTT AAAGTTCTGC TATGTGGCGC GGTATTATCC
CGTATTGACG CCGGGCAAGA GCAACTCGGT CGCCGCATAC ACTATTCTCA GAATGACTTG GTTGAGTACT CACCAGTCAC
AGAAAAGCAT CTTACGGATG GCATGACAGT AAGAGAATTA TGCAGTGCTG CCATAACCAT GAGTGATAAC ACTGCGGCCA
ACTTACTTCT GACAACGATC GGAGGACCGA AGGAGCTAAC CGCTTTTTTG CACAACATGG GGGATCATGT AACTCGCCTT
GATCGTTGGG AACCGGAGCT GAATGAAGCC ATACCAAACG ACGAGCGTGA CACCACGATG CCTGTAGCAA TGGCAACAAC
GTTGCGCAAA CTATTAACTG GCGAACTACT TACTCTAGCT TCCCGGCAAC AATTAATAGA CTGGATGGAG GCGGATAAAG
TTGCAGGACC ACTTCTGCGC TCGGCCCTTC CGGCTGGCTG GTTTATTGCT GATAAATCTG GAGCCGGTGA GCGTGGAAGC
CGCGGTATCA TTGCAGCACT GGGGCCAGAT GGTAAGCCCT CCCGTATCGT AGTTATCTAC ACGACGGGGA GTCAGGCAAC
TATGGATGAA CGAAATAGAC AGATCGCTGA GATAGGTGCC TCACTGATTA AGCATTGGTA A

**[6131–6950] Plasmid backbone (ori/noncoding)**

*Remaining backbone sequence including origin-associated/noncoding region*

CTGTCAGACC AAGTTTACTC ATATATACTT TAGATTGATT TAAAACTTCA TTTTTAATTT AAAAGGATCT AGGTGAAGAT
CCTTTTTGAT AATCTCATGA CCAAAATCCC TTAACGTGAG TTTTCGTTCC ACTGAGCGTC AGACCCCGTA GAAAAGATCA
AAGGATCTTC TTGAGATCCT TTTTTTCTGC GCGTAATCTG CTGCTTGCAA ACAAAAAAAC CACCGCTACC AGCGGTGGTT
TGTTTGCCGG ATCAAGAGCT ACCAACTCTT TTTCCGAAGG TAACTGGCTT CAGCAGAGCG CAGATACCAA ATACTGTTCT
TCTAGTGTAG CCGTAGTTAG GCCACCACTT CAAGAACTCT GTAGCACCGC CTACATACCT CGCTCTGCTA ATCCTGTTAC
CAGTGGCTGC TGCCAGTGGC GATAAGTCGT GTCTTACCGG GTTGGACTCA AGACGATAGT TACCGGATAA GGCGCAGCGG
TCGGGCTGAA CGGGGGGTTC GTGCACACAG CCCAGCTTGG AGCGAACGAC CTACACCGAA CTGAGATACC TACAGCGTGA
GCTATGAGAA AGCGCCACGC TTCCCGAAGG GAGAAAGGCG GACAGGTATC CGGTAAGCGG CAGGGTCGGA ACAGGAGAGC
GCACGAGGGA GCTTCCAGGG GGAAACGCCT GGTATCTTTA TAGTCCTGTC GGGTTTCGCC ACCTCTGACT TGAGCGTCGA
TTTTTGTGAT GCTCGTCAGG GGGGCGGAGC CTATGGAAAA ACGCCAGCAA CGCGGCCTTT TTACGGTTCC TGGCCTTTTG
CTGGCCTTTT GCTCACATGT

**Supplementary Table 3. Annotated full nucleotide sequence of pAAV-d′ (CMV-Region d)-mCherry**

| **Element** | **Coordinates (nt)** | **Length (bp)** | **Description** |
| --- | --- | --- | --- |
| 5′ AAV2 ITR | 1–145 | 145 | 5′ inverted terminal repeat |
| CMV enhancer | 146–530 | 385 | Human cytomegalovirus immediate-early enhancer sequence inserted upstream of Region d |
| XbaI (ITR/CMV enhancer junction) | 142–147 | 6 | Restriction site overlapping the 5′ ITR/CMV enhancer junction |
| Region d | 531–1143 | 613 | Candidate SLC2A1-associated regulatory element |
| XhoI + Kozak | 1144–1155 | 12 | Cloning site and Kozak sequence immediately upstream of mCherry |
| mCherry CDS | 1156–1860 | 705 | Coding sequence of mCherry, including stop codon |
| Spacer | 1861–2635 | 775 | Inserted spacer sequence between mCherry and WPRE |
| WPRE | 2636–3229 | 594 | Woodchuck hepatitis virus posttranscriptional regulatory element |
| Intervening noncoding region | 3230–3354 | 125 | Noncoding sequence between WPRE and the 3′ ITR |
| 3′ AAV2 ITR | 3355–3499 | 145 | 3′ inverted terminal repeat |
| Plasmid backbone | 3500–4531 | 1032 | Non-expression-cassette backbone sequence |
| bla / AmpR | 4532–5392 | 861 | Beta-lactamase coding sequence conferring ampicillin resistance |
| Plasmid backbone (ori/noncoding) | 5393–6212 | 820 | Remaining backbone sequence including origin-associated/noncoding region |

**Full nucleotide sequence**

**[1–145] 5′ AAV2 ITR**

*5′ inverted terminal repeat*

CCTGCAGGCA GCTGCGCGCT CGCTCGCTCA CTGAGGCCGC CCGGGCAAAG CCCGGGCGTC GGGCGACCTT TGGTCGCCCG GCCTCAGTGA GCGAGCGAGC

GCGCAGAGAG GGAGTGGCCA ACTCCATCAC TAGGGGTTCC TTCTA

**[146–530] CMV enhancer**

*Human cytomegalovirus immediate-early enhancer sequence*

GACTCGACAT TGATTATTGA CTAGTTATTA ATAGTAATCA ATTACGGGGT CATTAGTTCA TAGCCCATAT ATGGAGTTCC GCGTTACATA ACTTACGGTA

AATGGCCCGC CTGGCTGACC GCCCAACGAC CCCCGCCCAT TGACGTCAAT AATGACGTAT GTTCCCATAG TAACGCCAAT AGGGACTTTC CATTGACGTC

AATGGGTGGA GTATTTACGG TAAACTGCCC ACTTGGCAGT ACATCAAGTG TATCATATGC CAAGTACGCC CCCTATTGAC GTCAATGACG GTAAATGGCC

CGCCTGGCAT TATGCCCAGT ACATGACCTT ATGGGACTTT CCTACTTGGC AGTACATCTA CGTATTAGTC ATCGCTATTA CCATG

**[142–147] XbaI (ITR/CMV enhancer junction)**

*Restriction site overlapping the 5′ ITR/CMV enhancer junction*

TCTAGA

**[531–1143] Region d**

*Candidate SLC2A1-associated regulatory element*

CCTGGAGTCT CTCTAACAAA TAAATACTAC TTTCCATGCA GTAGACGCTG TTCTAAACAC TTTACAAATA TTAACTCACT TGGTCTTTCT ACAACCCTAC

GAGGTGGTAC TGTTACTATC CCTAGTGCAC CGAAGTCACC CAGCGGCCGA GTGAGAACTC CAGTCCAGCT TTCCACCCGC TACTCCGCGC ATCCCAGCTT

GCCTTACAGC CGGGTACCGG CTCCACCATT TTGCTAGAGA AGGCCGCGGA GGCTCAGAGA GGTGCGCACA CTTGCCCTGA GTCACACAGC GAATGCCCTC

CGCGGTCCCA ACGCAGAGAG AACGAGCCGA TCGGCAGCCT GAGCGAGGCA GTGGTTAGGG GGGGCCCCGG CCCCGGCCAC TCCCCTCACC CCCTCCCCGC

AGAGCGCCGC CCAGGACAGG CTGGGCCCCA GGCCCCGCCC CGAGGTCCTG CCCACACACC CCTGACACAC CGGCGTCGCC AGCCAATGGC CGGGGTCCTA

TAAACGCTAC GGTCCGCGCG CTCTCTGGCA AGAGGCAAGA GGTAGCAACA GCGAGCGTGC CGGTCGCTAG TCGCGGGTCC GCGAGTGAGC ACGCCAGGGA

GCAGGAGACC AAA

**[1144–1155] XhoI + Kozak**

*Cloning site and Kozak sequence immediately upstream of mCherry*

CTCGAGGCCA CC

**[1156–1860] mCherry CDS**

*Coding sequence of mCherry, including stop codon*

ATGGTGAGCA AGGGCGAGGA GGTCATCAAA GAGTTCATGC GCTTCAAGGT GCGCATGGAG GGCTCCATGA ACGGCCACGA GTTCGAGATC GAGGGCGAGG

GCGAGGGCCG CCCCTACGAG GGCACCCAGA CCGCCAAGCT GAAGGTGACC AAGGGCGGCC CCCTGCCCTT CGCCTGGGAC ATCCTGTCCC CCCAGTTCAT

GTACGGCTCC AAGGCGTACG TGAAGCACCC CGCCGACATC CCCGATTACA AGAAGCTGTC CTTCCCCGAG GGCTTCAAGT GGGAGCGCGT GATGAACTTC

GAGGACGGCG GTCTGGTGAC CGTGACCCAG GACTCCTCCC TGCAGGACGG CACGCTGATC TACAAGGTGA AGATGCGCGG CACCAACTTC CCCCCCGACG

GCCCCGTAAT GCAGAAGAAG ACCATGGGCT GGGAGGCCTC CACCGAGCGC CTGTACCCCC GCGACGGCGT GCTGAAGGGC GAGATCCACC AGGCCCTGAA

GCTGAAGGAC GGCGGCCACT ACCTGGTGGA GTTCAAGACC ATCTACATGG CCAAGAAGCC CGTGCAACTG CCCGGCTACT ACTACGTGGA CACCAAGCTG

GACATCACCT CCCACAACGA GGACTACACC ATCGTGGAAC AGTACGAGCG CTCCGAGGGC CGCCACCACC TGTTCCTGTA CGGCATGGAC GAGCTGTACA

AGTGA

**[1861–2635] Spacer**

*Inserted spacer sequence between mCherry and WPRE*

ACCCAGCTTT CTTGTACAAA GTGGGCATTA CTTATTGTTT TAGCTGTCCT CATGAATGTC TTTTCACTAC CCATTTGCTT ATCCTGCATC TCTCAGCCTT

GACTCCACTC AGTTCTCTTG CTTAGAGATA CCACCTTTCC CCTGAAGTGT TCCTTCCATG TTTTACGGCG AGATGGTTTC TCCTCGCCTG GGTCGAGCAG

TGTGGTTTTG CAAGAGGAAG CAAAAAGCCT CTCCACCCAG GCCTGGAATG TTTCCACCCA ATGTCGAGCA GTGTGGTTTT GCAAGAGGAA GCAAAAAGCC

TCTCCACCCA GGCCTGGAAT GTTTCCACCC AATGTCGAGC AAACCCCGCC CAGCGTCTTG TCATTGGCGA ATTCGAACAC GCAGATGCAG TCGGGGCGGC

GCGGTCCCAG GTCCACTTCG CATATTAAGG TGACGCGTGT GGCCTCGAAC ACCGAGCGAC CCTGCAGCCA ATATGGGATC GGCCATTGAA CAAGATGGAT

TGCACGCAGG TTCTCCGGCC GCTTGGGTGG AGAGGCTATT CGGCTATGAC TGGGCACAAC AGACAATCGG CTGCTCTGAT GCCGCCGTGT TCCGGCTGTC

AGCGCAGGGG CGCCCGGTTC TTTTTGTCAA GACCGACCTG TCCGGTGCCC TGAATGAACT GCAGGACGAG GCAGCGCGGC TATCGTGGCT GGCCACGACG

GGCGTTCCTT GCGCAGCTGT GCTCGACGTT GTCACTGAGC TCGAATTCGA TATCAAGCTT ATCGATAATT CCGAT

**[2636–3229] WPRE**

*Woodchuck hepatitis virus posttranscriptional regulatory element*

AATCAACCTC TGGATTACAA AATTTGTGAA AGATTGACTG GTATTCTTAA CTATGTTGCT CCTTTTACGC TATGTGGATA CGCTGCTTTA ATGCCTTTGT

ATCATGCTAT TGCTTCCCGT ATGGCTTTCA TTTTCTCCTC CTTGTATAAA TCCTGGTTGC TGTCTCTTTA TGAGGAGTTG TGGCCCGTTG TCAGGCAACG

TGGCGTGGTG TGCACTGTGT TTGCTGACGC AACCCCCACT GGTTGGGGCA TTGCCACCAC CTGTCAGCTC CTTTCCGGGA CTTTCGCTTT CCCCCTCCCT

ATTGCCACGG CGGAACTCAT CGCCGCCTGC CTTGCCCGCT GCTGGACAGG GGCTCGGCTG TTGGGCACTG ACAATTCCGT GGTGTTGTCG GGGAAGCTGA

CGTCCTTTCC ATGGCTGCTC GCCTGTGTTG CCACCTGGAT TCTGCGCGGG ACGTCCTTCT GCTACGTCCC TTCGGCCCTC AATCCAGCGG ACCTTCCTTC

CCGCGGCCTG CTGCCGGCTC TGCGGCCTCT TCCGCGTCTT CGCCTTCGCC CTCAGACGAG TCGGATCTCC CTTTGGGCCG CCTCCCCGCA TCGG

**[3230–3354] Intervening noncoding region**

*Noncoding sequence between WPRE and the 3′ ITR*

GAATTCCTAG AGCTCGCTGA TCAGCCTCGA CTGTGCCTTC TAGTTGCCAG CCATCTGTTG TTTGCCCCTC CCCCGTGCCT TCCTTGACCC TGGAAGGTGC

CACTCCCACT GTCCTTTCCT AATAA

**[3355–3499] 3′ AAV2 ITR**

*3′ inverted terminal repeat*

AATGAGGAAA TTGCATCGCA TTGTCTGAGT AGGTGTCATT CTATTCTGGG GGGTGGGGTG GGGCAGGACA GCAAGGGGGA GGATTGGGAA GAGAATAGCA

GGCATGCTGG GGAGGGCCGC AGGAACCCCT AGTGATGGAG TTGGC

**[3500–4531] Plasmid backbone**

*Non-expression-cassette backbone sequence*

CACTCCCTCT CTGCGCGCTC GCTCGCTCAC TGAGGCCGGG CGACCAAAGG TCGCCCGACG CCCGGGCTTT GCCCGGGCGG CCTCAGTGAG CGAGCGAGCG

CGCAGCTGCC TGCAGGGGCG CCTGATGCGG TATTTTCTCC TTACGCATCT GTGCGGTATT TCACACCGCA TACGTCAAAG CAACCATAGT ACGCGCCCTG

TAGCGGCGCA TTAAGCGCGG CGGGGGTGGT GGTTACGCGC AGCGTGACCG CTACACTTGC CAGCGCCTTA GCGCCCGCTC CTTTCGCTTT CTTCCCTTCC

TTTCTCGCCA CGTTCGCCGG CTTTCCCCGT CAAGCTCTAA ATCGGGGGCT CCCTTTAGGG TTCCGATTTA GTGCTTTACG GCACCTCGAC CCCAAAAAAC

TTGATTTGGG TGATGGTTCA CGTAGTGGGC CATCGCCCTG ATAGACGGTT TTTCGCCCTT TGACGTTGGA GTCCACGTTC TTTAATAGTG GACTCTTGTT

CCAAACTGGA ACAACACTCA ACTCTATCTC GGGCTATTCT TTTGATTTAT AAGGGATTTT GCCGATTTCG GTCTATTGGT TAAAAAATGA GCTGATTTAA

CAAAAATTTA ACGCGAATTT TAACAAAATA TTAACGTTTA CAATTTTATG GTGCACTCTC AGTACAATCT GCTCTGATGC CGCATAGTTA AGCCAGCCCC

GACACCCGCC AACACCCGCT GACGCGCCCT GACGGGCTTG TCTGCTCCCG GCATCCGCTT ACAGACAAGC TGTGACCGTC TCCGGGAGCT GCATGTGTCA

GAGGTTTTCA CCGTCATCAC CGAAACGCGC GAGACGAAAG GGCCTCGTGA TACGCCTATT TTTATAGGTT AATGTCATGA TAATAATGGT TTCTTAGACG

TCAGGTGGCA CTTTTCGGGG AAATGTGCGC GGAACCCCTA TTTGTTTATT TTTCTAAATA CATTCAAATA TGTATCCGCT CATGAGACAA TAACCCTGAT

AAATGCTTCA ATAATATTGA AAAAGGAAGA GT

**[4532–5392] bla / AmpR**

*Beta-lactamase coding sequence conferring ampicillin resistance*

ATGAGTATTC AACATTTCCG TGTCGCCCTT ATTCCCTTTT TTGCGGCATT TTGCCTTCCT GTTTTTGCTC ACCCAGAAAC GCTGGTGAAA GTAAAAGATG

CTGAAGATCA GTTGGGTGCA CGAGTGGGTT ACATCGAACT GGATCTCAAC AGCGGTAAGA TCCTTGAGAG TTTTCGCCCC GAAGAACGTT TTCCAATGAT

GAGCACTTTT AAAGTTCTGC TATGTGGCGC GGTATTATCC CGTATTGACG CCGGGCAAGA GCAACTCGGT CGCCGCATAC ACTATTCTCA GAATGACTTG

GTTGAGTACT CACCAGTCAC AGAAAAGCAT CTTACGGATG GCATGACAGT AAGAGAATTA TGCAGTGCTG CCATAACCAT GAGTGATAAC ACTGCGGCCA

ACTTACTTCT GACAACGATC GGAGGACCGA AGGAGCTAAC CGCTTTTTTG CACAACATGG GGGATCATGT AACTCGCCTT GATCGTTGGG AACCGGAGCT

GAATGAAGCC ATACCAAACG ACGAGCGTGA CACCACGATG CCTGTAGCAA TGGCAACAAC GTTGCGCAAA CTATTAACTG GCGAACTACT TACTCTAGCT

TCCCGGCAAC AATTAATAGA CTGGATGGAG GCGGATAAAG TTGCAGGACC ACTTCTGCGC TCGGCCCTTC CGGCTGGCTG GTTTATTGCT GATAAATCTG

GAGCCGGTGA GCGTGGAAGC CGCGGTATCA TTGCAGCACT GGGGCCAGAT GGTAAGCCCT CCCGTATCGT AGTTATCTAC ACGACGGGGA GTCAGGCAAC

TATGGATGAA CGAAATAGAC AGATCGCTGA GATAGGTGCC TCACTGATTA AGCATTGGTA A

**[5393–6212] Plasmid backbone (ori/noncoding)**

*Remaining backbone sequence including origin-associated/noncoding region*

CTGTCAGACC AAGTTTACTC ATATATACTT TAGATTGATT TAAAACTTCA TTTTTAATTT AAAAGGATCT AGGTGAAGAT CCTTTTTGAT AATCTCATGA

CCAAAATCCC TTAACGTGAG TTTTCGTTCC ACTGAGCGTC AGACCCCGTA GAAAAGATCA AAGGATCTTC TTGAGATCCT TTTTTTCTGC GCGTAATCTG

CTGCTTGCAA ACAAAAAAAC CACCGCTACC AGCGGTGGTT TGTTTGCCGG ATCAAGAGCT ACCAACTCTT TTTCCGAAGG TAACTGGCTT CAGCAGAGCG

CAGATACCAA ATACTGTTCT TCTAGTGTAG CCGTAGTTAG GCCACCACTT CAAGAACTCT GTAGCACCGC CTACATACCT CGCTCTGCTA ATCCTGTTAC

CAGTGGCTGC TGCCAGTGGC GATAAGTCGT GTCTTACCGG GTTGGACTCA AGACGATAGT TACCGGATAA GGCGCAGCGG TCGGGCTGAA CGGGGGGTTC

GTGCACACAG CCCAGCTTGG AGCGAACGAC CTACACCGAA CTGAGATACC TACAGCGTGA GCTATGAGAA AGCGCCACGC TTCCCGAAGG GAGAAAGGCG

GACAGGTATC CGGTAAGCGG CAGGGTCGGA ACAGGAGAGC GCACGAGGGA GCTTCCAGGG GGAAACGCCT GGTATCTTTA TAGTCCTGTC GGGTTTCGCC

ACCTCTGACT TGAGCGTCGA TTTTTGTGAT GCTCGTCAGG GGGGCGGAGC CTATGGAAAA ACGCCAGCAA CGCGGCCTTT TTACGGTTCC TGGCCTTTTG

CTGGCCTTTT GCTCACATGT

**Supplementary Table 4 | Primers used for reporter plasmid construction and ATG elimination**

| **Category** | **Primer name** | **Sequence (5′–3′)** |
| --- | --- | --- |
| Reference (Ref.) | Reference_enhancer_Fw | aggggttccttctagaaccattttgctagagaagg |
| Reference (Ref.) | Reference_enhancer_Rv | ctcaccatggtggcctcgagagagagcgcgcggac |
| ATG elimination | hGLUT1_deltaATG_Fw | gccggggtcctataaacgctacggtccg |
| ATG elimination | hGLUT1_deltaATG_Rv | tggctggcgacgccggtgtgtc |
| Region a–c | Region_abc_Fw | cactaggggttccttctagatttgctagagaaggccg |
| Region a | Region_a_Rv | ccatggtggcctcgagcggacccgcgactagcgac |
| Region b | Region_b_Rv | ccatggtggcctcgagcaaagctttggggcctgca |
| Region c | Region_c_Rv | tggtggcctcgagccccttctgcccaaaattcaag |
| Region d | Region_d_Fw | atcactaggggttccttctagacctggagtctctc |
| Region d | Region_d_Rv | ctcaccatggtggcctcgagtttggtctcctgctc |
| Region d′ (for CMV） | Region_dprime_Fw | aggggttccttctagactcgacattgattattgac |
| Region d′ (for CMV） | Region_dprime_Rv | gttagagagactccaggcatggtaatagcgatgac |
| Region e | Region_e_Fw | cactaggggttccttctagaacagtgagccgagat |
| Region e | Region_e_Rv | cggccttctctagcaaaatggtggagccggtac |
| Region f–h | Region_fgh_Rv | tgctcaccatggtggcctcgagctcactcggggac |
| Region f | Region_f_Fw | cactaggggttccttctagagccttccctcaatcc |
| Region g | Region_g_Fw | cactaggggttccttctagattgaagcaaaatatg |
| Region h | Region_h_Fw | ggggttccttctagaaaagtctggggacaaaccag |
| Region i | Region_i_Fw | ctcaccatggtggcctcgaggcccaaaattcaaga |
| Region i | Region_i_Rv | cactaggggttccttctagaaaagtctggggacaa |

*Fw, forward primer; Rv, reverse primer. Identical primer sequences used for multiple regulatory elements are grouped under a shared primer name (for example, Region_abc_Fw and Region_fgh_Rv).*
